# Crunch: Integrated processing and modeling of ChIP-seq data in terms of regulatory motifs

**DOI:** 10.1101/042903

**Authors:** Severin Berger, Saeed Omidi, Mikhail Pachkov, Phil Arnold, Nicholas Kelley, Silvia Salatino, Erik van Nimwegen

## Abstract

Although it has become routine for experimental groups to apply ChIP-seq technology to quantitatively characterize the genome-wide binding of transcription factors (TFs), computational analysis procedures remain far from standardized, making it difficult to meaningfully compare ChIP-seq results across experiments. In addition, while genome-wide binding patterns must ultimately be determined by local constellations of binding sites in the DNA, current analysis is typically limited to a standard search for enriched motifs in ChIP-seq peaks.

Here we present Crunch, a completely automated computational method that performs all ChIP-seq analysis from quality control through read mapping and peak detecting, and integrates comprehensive modeling of the ChIP signal in terms of known and novel binding motifs, quantifying the contribution of each motif, and annotating which combinations of motifs explain each binding peak.

Applying Crunch to 128 ChIP-seq datasets from the ENCODE project we find that TFs naturally separate into ‘solitary TFs’, for which a single motif explains the ChIP-peaks, and ‘co-binding TFs’ for which multiple motifs co-occur within peaks. Moreover, for most datasets the motifs that Crunch identified *de novo* outperform known motifs and both the set of co-binding motifs and the top motif of solitary TFs are consistent across experiments and cell lines. Crunch is implemented as a web server (crunch.unibas.ch), enabling standardized analysis of any collection of ChIP-seq datasets by simply uploading raw sequencing data. Results are provided both in a graphical interface and as downloadable files.

## 3 Introduction

The advent of next-generation sequencing technologies and the associated dramatic reduction of cost for sequencing have led to a spectacular rise in the use of a variety of methods, including RNA-seq, ChIP-seq, DNase-seq, ATAC-seq and CLIP-seq, that combine next-generation sequencing with other molecular biology techniques to quantitatively characterize internal states of cells on a genome-wide scale (Buenrostro et al. 2013; Soon et al. 2013). As one of the most prominent of these technologies, ChIP-seq (Johnson et al. 2007) combines chromatin immuno-precipitation with next-generation sequencing to quantify the genome-wide binding patterns of any molecule that associates with the DNA. Apart from large-scale efforts, such as the ENCODE project in which ChIP-seq was used to systematically map the binding patterns of many transcription factors (TFs) (Bernstein et al. 2012), many individual labs are now using ChIP-seq to characterize the binding patterns of particular DNA-binding proteins of interest in their specific system of interest, e.g. particular tissues or specific biological conditions.

The result of a ChIP-seq experiment is a collection of short DNA sequence reads, i.e. typically tens of million reads that are a few tens of basepairs long each. Like for other high-throughput experimental techniques, extracting meaningful biological information from such large data-sets is a non-trivial computational task that involves a significant number of separate steps including read quality control and pre-processing, mapping the reads to the genome, estimating the typical length of the DNA fragments from which the reads derive, identifying binding peaks, i.e. identifying genomic regions that are statistically significantly enriched for immuno-precipitated fragments, and downstream analyses such as identification of sequence motifs enriched within the peak sequences. Over the last decade a large number of bioinformatic tools has been developed to perform each of these tasks, e.g. see (Schbath et al. 2012) for a review of read mapping tools, (Wilbanks and Facciotti 2010) for a review of methods for detecting binding peaks, and (Das and Dai 2007) for an overview of algorithms for finding sequence motifs over-represented among a set of short sequence segments. A number of tools have also been presented that allow researchers to combine individual tools into a workflow, i.e. by allowing users to manually execute one tool after another or by constructing a pipeline that runs the tools automatically. These include commercial solutions such as Avadis NGS (Strand Life Sciences Pvt. Ltd. 2012), Chipster (Kallio et al. 2011), CLCbio Genomics Workbench (CLC Bio 2017), Genomatix Mining Station (Genomatix 2012) and Partek Genomics Suite (Partek Inc. 2008), as well as free-to-use solutions such as HOMER (Heinz et al. 2010), cisGenome (Ji et al. 2008), seqMINER (Ye et al. 2011), ChIPseeqer (Giannopoulou and Elemento 2011), GeneProf (Halbritter et al. 2013) and Galaxy/Cistrome (Liu et al. 2011).

However, current ChIP-seq analysis methods are highly unsatisfactory in a number of re spects. First, there is no consensus on what tools are most appropriate for each step in the ChIP-seq analysis, let alone regarding details of their parameters and implementation, and this leads to a very large variability in the way ChIP-seq data is currently analyzed. The main challenge, especially for experimental researchers that wish to perform ChIP-seq analysis but lack computational biology expertise, is that there are no standardized pipe-lines that are widely accepted to give satisfactory performance. A private survey that we performed among colleagues suggests that every group with expertise in processing ChIP-seq data uses a different combination of tools that they have individually customized to deal with various issues that are not addressed by publically available tools. A consequence of this situation is that it is extremely difficult to compare results of ChIP-seq data from different experiments. That is, in order to compare one’s own results with those of other ChIP-seq experiments, it is nessecary to re-analyze the raw data from these experiments using one’s own customized analysis pipeline.

Another unsatisfactory aspect of current analysis practices is that popular peak finders, e.g. MACS (Zhang et al. 2008), assume statistical models for the fluctuations in ChIP-seq read densities that do not match those that are observed in the data. Consequently, the interpretation of the enrichment statistics (i.e. *p*-values) that such methods provide is problematic, and other methods, such as analysis of the reproducibility of peaks across replicates (Landt et al. 2012), are necessary to determine cut-offs on the statistical significance of peaks.

Probably the most unsatisfactory aspect of current ChIP-seq analysis practices concerns the analysis of DNA sequence motifs. Although the entire genome-wide ChIP signal should ultimately be determined by local constellations of binding sites for DNA-binding factors, current ChIP-seq analyses do not attempt to directly model the observed ChIP signal in terms of the underlying DNA sequence. Instead, downstream sequence analysis is typically limited to simply running standard motif finding algorithms on the sequences of the top binding peaks.

To address all these issues, we here present Crunch, a completely automated procedure that performs all steps of the ChIP-seq analysis and integrates comprehensive modeling of the genome-wide ChIP signal in terms of binding sites for both known and novel motifs, which Crunch identifies *de novo.* Crunch is implemented as a webserver at crunch.unibas.ch and only requires upload of the raw sequencing reads, allowing any researcher to perform comprehensive ChIP-seq analysis of any number of datasets in a completely standardized manner. Besides flat files for download, all results of Crunch’s analysis are available through an easily navigable graphical web interface. Among Crunchs novel features are a Bayesian mixture model that identifies significantly enriched regions by fitting a noise model directly to the data of each experiment, as well as a Gaussian mixture model for decomposing enriched regions into individual binding peaks. In addition, Crunch uses a combination of *de novo* motif finding with binding site prediction for a large collection of known regulatory motifs to model the observed ChIP-seq signal in terms of novel and known regulatory motifs, extensively characterizing the contribution of each motif to explaining the ChIP-seq signal, and annotating which combinations of motifs occur in each binding peak.

To demonstrate the power of the method we apply Crunch to a large collection of ChIP-seq datasets from the ENCODE project. We show that the motifs that Crunch identifies *de novo* often outperform all known motifs and that Crunch successfully identifies binding partners of the proteins that were immuno-precipitated. Interestingly, we find that TFs divide into ‘solitary binding TFs’ where the observed peaks are explained by a single underlying sequence motif, and ‘co-binding TFs’ where multiple motifs, often including known motifs for other TFs, are necessary to explain the observed binding peaks. Moreover, the set of co-binding motifs are highly reproducible across different ChIP-seq experiments done on different cell lines. Similarly, we find that the top motif for solitary binding TFs is also highly reproducible across experiments, even when very similar motifs from different collections are available, suggesting that Crunch can be used to identify the optimal *in vivo* binding motif for a given solitary TF.

## 4 Results

In a typical ChIP-seq experiment the protein that is immuno-precipitated is a DNA-binding protein, or a protein that forms complexes with DNA-binding proteins, and the aims of the experiment include identifying the genomic loci where the protein is binding (either directly or indirectly), and the genes that are potentially regulated by these binding events. For DNA-binding proteins that bind DNA in a sequence-specific manner, additional aims are to characterize the sequence specificity of the protein, and also to identify other DNA-binding proteins that are co-localizing with the immuno-precipitated factor, possibly through direct interactions. Crunch provides answers to all of these questions and makes its analysis results accessible through an interactive graphical web interface and downloadable files. In addition, Crunch provides extensive reports on all steps of the analysis that allow researchers to assess the quality of the data and to troubleshoot potential problems with the data.

To make the Crunch analysis easily available to any researcher, including those without computational biology expertise, the webserver at crunch.unibas requires only a minimal amount of information from the user. All the user needs to provide is raw ChIP-seq data in the form of FASTQ or FASTA files, and to provide the organism from which the data derived (currently human, mouse, and Drosophila have been implemented). Although a single immuno-precipitation dataset, i.e. a ‘foreground’, suffices, users are advised to also upload corresponding ‘background’ samples, i.e. input DNA, whenever these are available. Multiple replicate datasets can either be uploaded and analyzed separately or analyzed jointly. Optionally, more advanced users can choose to edit a number of options (see Methods) including the possibility to upload BED files of already mapped data instead of raw sequence data. Users can specify an e-mail address to get an automatic notification with a link to the results when the analysis has finished.

We used Crunch to analyse a large set of ChIP-seq experiments from the ENCODE consortium (Bernstein et al. 2012). We chose to analyze all experiments performed on the cell line GM12878 and to additionally all experiments that were performed on the HeLaS3 cell line by the laboratory of Michael Snyder at Stanford University. In total we analysed 128 experiments in which 93 different factors were immuno-precipitated. The full reports that result from submitting the raw FASTQ data of all these ChIP-seq datasets to Crunch, as well as links to the raw data, are available at crunch.unibas.ch/ENCODE_REPORTS. Below we will first use a single ChIP-seq dataset from this collection to discuss all steps of the Crunch analysis as well as illustrate the results that each of these analyses provide. After that, we will discuss a number of new insights that result from the results of Crunch’s analysis on the full collection of ChIP-seq datasets.

### Analysis and report overview

To illustrate the results that Crunch provides, we chose an experiment from the ENCODE collection where the BRCA1 protein was immunoprecipitated from GM12878 cells. Two replicate foreground (immuno-precipitation) samples as well as two replicate background (input DNA) samples were produced.

The analysis performed by Crunch is structured into three parts, as schematized in Fig. 1, and the analysis report is structured accordingly. In the first pre-processing part, the reads from each submitted sample are analyzed and filtered for quality, they are mapped to the genome, and the mappings are used to estimate the sizes of the DNA fragments in the sample. In the second ‘peak calling’ part a Bayesian mixture model is fitted to identify genomic regions that are significantly enriched in the ChIP samples, these regions are then further separated into individual binding peaks using a Gaussian mixture model, and the final peaks are annotated with locations of nearby genes. In the third ‘motif analysis’ part, novel binding motifs are inferred *de novo* and the peak sequences are then modeled in terms of these novel and a library of known regulatory motifs. We now discuss each of these parts in turn for the BRCA1 example. Detailed descriptions of all analysis steps are provided in the Methods section.

**Figure 1:**
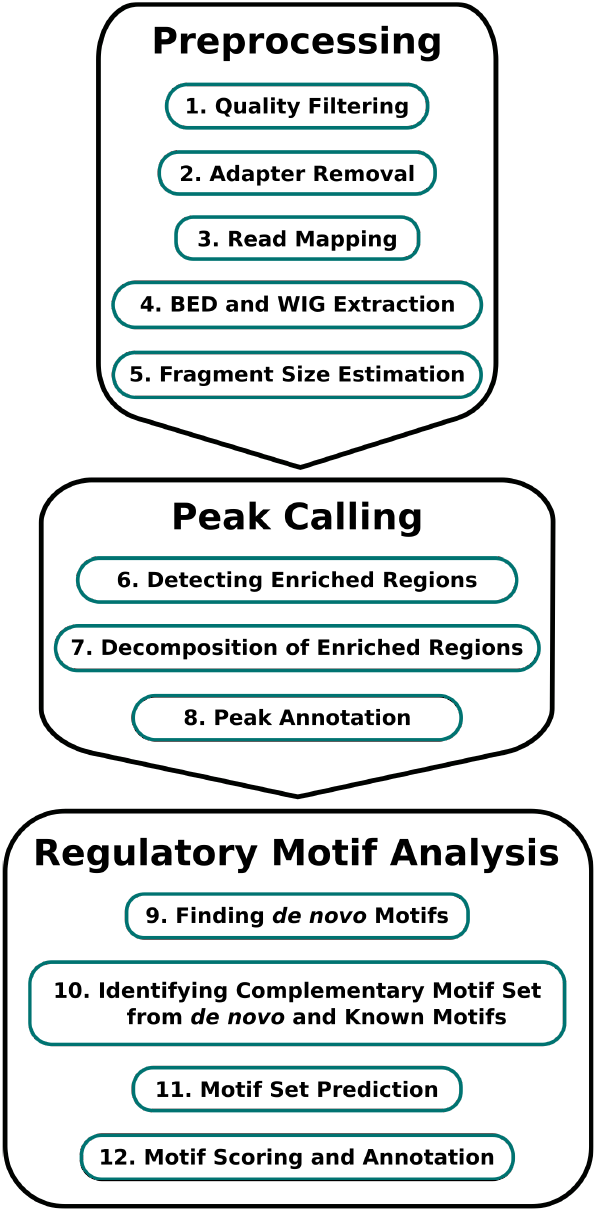
An overview of the steps in Crunch’s ChIP-seq analysis. The analysis can be divided into three parts: pre-processing of the data, identification of the binding peaks, and regulatory motif analysis.

### Quality Control, Mapping, and Fragment Size Estimation

The quality control (QC), mapping, and fragment length estimation are performed separately for each submitted sample. The quality control consists of first removing reads of low sequence quality. Next, Crunch uses a library of known 3’ end adapters to search for adapter sequences that still remain at the 5’ ends of reads and trims these adapter sequences from the reads. Finally, reads with low dinucleotide complexity are removed. These QC steps are fairly standard and, for any dataset of reasonable quality, only a small fraction of the reads should be removed in this step, i.e. for the first BRCA1 ChIP sample almost 93% of the reads pass this filter (Fig. 2).

**Figure 2:**
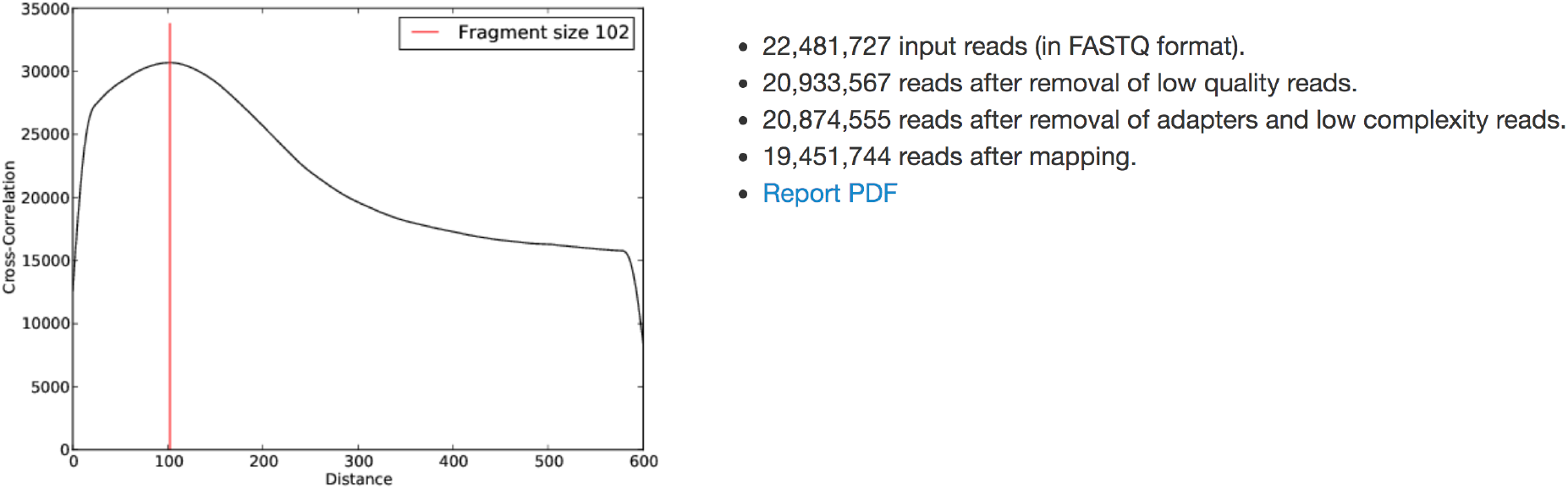
Summary report of the quality control, mapping and fragment size estimation for the first ChIP sample (foreground) of the BRCA1 dataset from the GM12878 cell line. On the right the numbers of reads passing various steps of the quality control and mapping are shown. On the left the cross-correlation function *C*(*d*) of co-occurence of reads mapping to opposite strands at a distance *d* is shown. The typical fragment length is estimated as the maximum of the cross-correlation function (red line).

Crunch’s read mapping procedure is also relatively standard and performed using Bowtie (Langmead et al. 2009). Although in some approaches only uniquely mapping reads are maintained, we feel that this is problematic because this will lead to a systematic loss of binding peaks in regions whose sequence is not unique in the genome. We thus retain multi-mapping reads but equally distribute the weight of each read to all mapping locations (see Methods). For the first BRCA1 ChIP sample, a little over 93% of the reads are successfully mapped to the genome. Loss of a large number of reads at any of the QC or mapping steps may indicate low data quality. A more extensive report with detailed statistics regarding the quality control, adapter removal and mappings is provided as a PDF for download (see Suppl. Fig. S1). For each sample, Crunch also provides a downloadable BED file with the mappings, and a WIG file that allows visualization of the read density along the genome.

In most ChIP-seq protocols the sequenced reads derive from longer DNA fragments and, for the downstream analysis, it is important to estimate the typical size of these fragments. Since each read is in principle equally likely to derive from either the 5’ or 3’ end of a fragment, each fragment of length *l* leads to pairs of reads on opposite strands at a distance *l*. Consequently, when the typical fragment length is *l*, co-occurring reads on opposite strands should be more common at distance *l* than at other distances (Schmid and Bucher 2007). To estimate the fragment length we calculate, for each distance *d*, the number of genomic positions *C*(*d*) for which reads are observed on both strands at a distance *d* from each other. We estimate the typical fragment length l as the value of *d* for which *C*(*d*) is maximal (Fig. 2, Suppl. Fig. S2, and Methods). In most cases the correlation function *C*(*d*) shows a clear peak indicating that Crunch could unambiguously infer the fragment size. However, if no definite peak is visible or the estimated fragment size does not clearly match a peak in the profile, this indicates that there was no typical fragment length, which in turn may indicate that the sample had low quality.

### Binding peak identification and annotation

The aim of the peak identification part of the analysis is to find all locations in the genome where the density of fragments in the ChIP sample was significantly higher than in the background (i.e. input DNA) sample. To do this, we need to estimate the density of ChIP fragments in both the ChIP and background samples from the mapped reads, and develop a statistical model for the fluctuations in these densities in order to determine which genomic regions are significantly enriched.

Crunch first calculates the observed read densities in sliding windows along the genome for both the ChIP and background samples (see Methods). Although in principle the distribution of read densities in the background sample is expected to be roughly constant, we observe that there is a small fraction of windows, typically around 0.1%, and typically associated with genomic repeats, where read densities in the background sample are much higher than expected (Fig. 3A). These regions might correspond to repeats that are more common in the genomes of the cells used in the experiment than in the reference assembly. Since these regions show aberrant read densities in both ChIP and background samples, statistically significant enrichment of ChIP reads cannot be reliably assessed in these regions, and Crunch uses an adaptive procedure to filter out this small subset of windows (see Methods).

**Figure 3:**
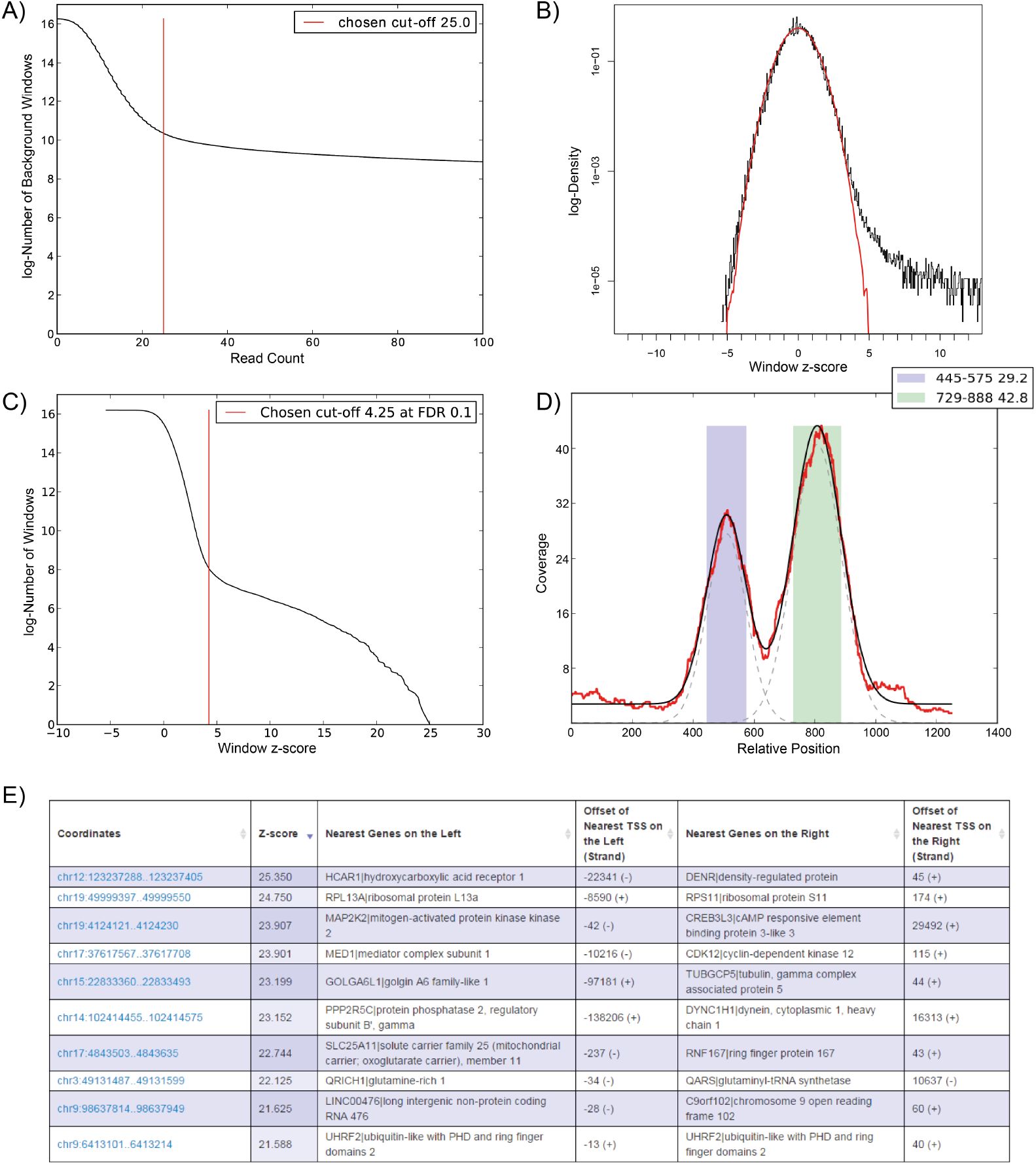
A) Reverse cumulative distribution of the summed read counts from the BRCA1 background samples in genome-wide sliding windows of 2000 bps. Crunch adaptively determines a cut-off (red line) and windows with coverage higher than this cut-off are removed from further analysis. B) Distribution of the observed *z*-values from all genome-wide sliding widows (in black) and a reference standard normal distribution (in red). C) Reverse cumulative distribution of the same z-values (in black) as well as the z-score threshold (in red) corresponding to an FDR of 0.1. Note that the vertical axes in panels A, B and C are in log-scale. D) ChIP read density profile of an individual enriched region (red), together with the fitted mixture model (black). The two grey dashed lines show the Gaussians used in the mixture model of this region. The coloured bars show the two individual binding peaks with their locations and amplitudes shown in the legend. E) Table with the top 10 peaks of the BRCA1 experiment. Each peak is annotated with its coordinates on the genome, its z-value, its nearest up- and down-stream genes and the distance to the transcription start sites of these genes.

To detect windows that are significantly enriched in the ChIP sample relative to the background, we need a statistical model for the fluctuations in observed read densities for windows that are *not* enriched. It is well known that, even for identical sequence libraries, the act of sequencing itself introduces Poisson sampling noise in the number of observed reads in a given window. While some peak finders make the assumption that this Poisson sampling noise is the *only* source of fluctuations, it is also well recognized that the other steps in the protocol such as the fragmenting of the DNA and PCR amplification introduce additional noise. For reasons of mathematical convenience, it has become popular to assume that these additional fluctuations are Gamma distributed, leading to an overall negative binomial distribution of read densities across replicates, see e.g. (Anders and Huber 2010). However, in previous work we have shown that the empirically observed fluctuations in read densities across replicates are well described by a convolution of multiplicative, i.e. log-normal, and Poisson sampling noise (Balwierz et al. 2009).

As detailed in the methods, we thus developed a Bayesian mixture model which assumes that the genomic windows derive from a mixture of unenriched and enriched regions. In regions that are unenriched, the difference between ChIP and background read densities is assumed to follow a ‘background distribution’ that can be described as a convolution of log-normal and Poisson sampling noise, while in the enriched regions the ChIP read density can be arbitrarily higher than in the background read density. The Bayesian mixture model fits the parameters of the background distribution as well as the total fraction of enriched windows and calculates, for each window, a posterior probability that the window is enriched. This posterior probability can be easily expressed in terms of a *z*-statistic, which gives the number of standard-deviations by which the ChIP density is higher than the background density (see Methods). Crunch’s noise model predicts that, in the absence of any enriched windows, the *z*-values of windows across the genome should follow a standard normal distribution. As shown in Fig. 3B, for the BRCA1 dataset we indeed observe that the *z*-statistics of more than 99.9% of the windows almost perfectly follow a standard-normal distribution, confirming that the data support the noise model that Crunch assumes. Figure 3B also shows a small tail of windows with much higher *z*-values, corresponding to significantly enriched windows. To select the set of significantly enriched windows Crunch orders windows by their *z*-value (or posterior probability to be enriched) and picks a cut-off such that the expected fraction of selected windows that are not truly enriched, i.e. the false discovery rate (FDR), is equal to 0.1 (Fig. 3C).

Crunch then merges enriched windows that overlap on the genome into enriched genomic regions. As shown in Fig. 3D, the ChIP read density profiles in a single enriched genomic region may show evidence of multiple individual binding peaks. Theoretically, when the immuno-precipitated protein is bound to the genome at a single location, this should lead to a peak in the ChIP signal, centered at this location, with a width determined by the typical length of the immuno-precipitated fragments and this is confirmed by our analysis of the ENCODE ChIP-seq datasets (see Methods and Suppl. Text S1). Using this observation, Crunch separates each genomic region into one or more binding peaks by modeling the observed ChIP read density profile as a mixture of a constant background plus a collection of Gaussians, with widths constrained by the estimated typical fragment length (see Methods and Fig. 3D). A *z*-value is recomputed for each individual peak, and only those peaks are retained that pass the previously determined *z*-value cut-off (indicated as the green and purple segments in Fig. 3D). Finally, the results of the peak finding are summarized as a list with genomic coordinates, *z*-value, and the locations of the nearest transcription start sites of genes up- and down-stream of each binding peak (Fig. 3E).

### Explaining the observed peak sequences by a complementary set of motifs

Probably the most novel aspect of Crunch is that it aims to explicitly explain the observed peak sequences in terms of sequence motifs that occur within them. In order to make this notion precise, we employ an idealized model of the ChIP-seq experiment. We assume that the probability for a particular genomic sequence to end up in the list of binding peaks is proportional to the total amount of binding of the immuno-precipitated protein to this sequence. Next, we assume that the immuno-precipitated protein associates with a particular set of DNA-binding factors, whose sequence-specificities can be represented by a set of motifs {*w*}, i.e. a set of position specific weight matrices (PWMs). In particular, we assume that the probability *P*(*s*) that a particular sequence s ends up as a binding peak is proportional to the total number of binding sites *n*_{*w*}_ (*s*) in sequence s for the motifs of the set {*w*}, i.e. *P*(*s*) ∝ *n*_{*w*}_(*s*). Given a set of motifs {*w*} and the set of observed peak sequences *S*, we imagine this set *S* embedded in a large pool of randomized sequences with the same nucleotide composition, and then calculate the probability *P*(*S*|{*w*}) of observing *only* the set of sequences S as binding peaks, and none of the randomized sequences. Crunch then searches for an optimal set of motifs {*w*} that maximizes this probability *P*(*S*|{*w*}), using a library of known motifs, as well as motifs that were identified *de novo*.

Crunch first selects all binding peak sequences and randomly divides these into a training set and a test set of equal size (if there are more than 1000 binding peaks only the top 1000 peaks are used for computational efficiency). Crunch then first performs *de novo* motif finding on the training set using the PhyloGibss (Siddharthan et al. 2005) and MotEvo (Arnold et al. 2012) algorithms previously developed in our labs. Both these algorithms are designed to incorporate information from sequence conservation patterns by running on multiple alignments of orthologous genomic regions. Crunch automatically aligns each peak sequence in the training set with orthologous sequences from related organisms and runs PhyloGibbs and MotEvo on these multiple alignments (see Methods).

The motifs that were found *de novo* are then combined with a large collection of over 2000 PWMs that we collected from a number of resources into a motif library *W* (see Methods). The training and test sets of peak sequences are then augmented with a large pool of randomized sequences that have identical nucleotide composition as the peak sequences. Using MotEvo, we then iteratively add motifs from our library *W* to a set of selected motifs {*w*} so as to maximize the probability *P*(*S*|{*w*}) for the sequences *S* of the test set, until addition of any motif to the set {*w*} no longer significantly increases the probability *P*(*S*|{*w*}) (see Methods). The result is a set of complementary motifs that jointly best explain the observed binding peak sequences *S* (Fig. 4A).

**Figure 4:**
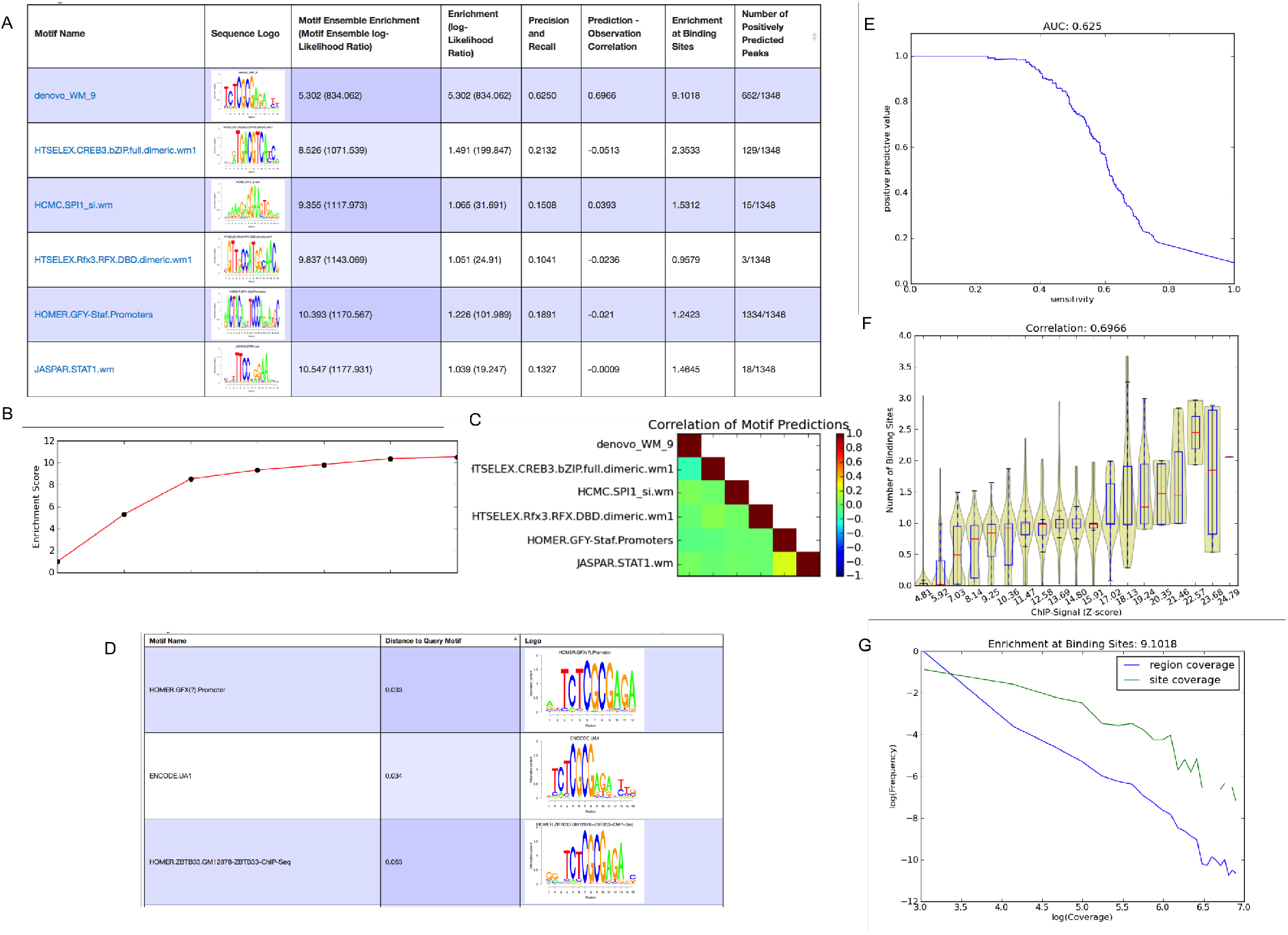
A) List of complementary motifs that jointly explain the BRCA1 binding peaks. The motifs are sorted in order of their contribution to explaining the peaks and for each motif its name, sequence logo, and a set of statistics characterizing the performance of the motif are shown. B) Evolution of the enrichment score of the motif set {*w*} as more motifs are added. C) Heatmap of pairwise correlations of the occurrence of all motifs in {*w*} across binding peaks. D) The top three motifs from the library of known motifs with most similarity to the novel motif denovo_WM_9. E) Precision-recall curve for motif denovo_WM_9. The area under the curve (AUC) is 0.625. F) Correlation between the ChIP signal (peak z-scores) and the number of predicted binding sites for denovo_WM_9 within peaks. The Pearson correlation is 0.6966. G) Distributions of ChIP-coverage at denovo_WM_9 sites (green curve) and at all positions in peaks (blue curve). The coverage at sites is on average 9.1018 as high as at random positions in the peaks.

To quantify the quality of a motif set {*w*} with an intuitive measure, Crunch calculates the enrichment score 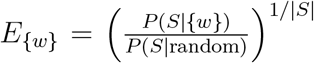, where *P*(*S*|random) is the probability to select the binding peak sequences S by chance, and |*S*| is the number of binding peaks. The enrichment score quantifies how much more likely, on average, the peak sequences are to be selected given the occurrence of the motifs {*w*} compared to when selecting sequences at random. Crunch reports, for each motif, its enrichment score when considered by itself (Fig. 4A). For the BRCA1 peaks the top motif was a novel motif found de *novo,* called denovo_WM_9, with an enrichment score of about 5.3. That is, the peak sequences on average have 5.3 times as many sites for this motif as random sequences of the same nucleotide composition. Note also that the enrichment score for all other motifs in the set is below 1.5. However, when these motifs are combined with denovo_WM_9, the enrichment increases as shown in the first column of the table in Fig. 4A and in Fig. 4B. When all 6 motifs are considered jointly, the enrichment score has almost doubled to 10.547. Crunch also investigates to what extent different motifs in the set tend to either co-occur or avoid co-occurring within the same peak sequences, and reports the correlations in motif occurrence for all pairs of motifs in the set (Fig. 4C).

For each motif in the set {*w*}, an extensive set of additional information and performance measures are reported. First, besides showing the motif’s sequence logo, each motif is compared with all other motifs in the library of known motifs, and a list of the top matching motifs is reported (Fig. 4D). We see that Crunch’s top motif for the BRCA1 data, i.e. denovo_WM_9, is similar to a GFX motif in Homer, the UA1 motif from ENCODE, and a motif for the TF ZBTB33 from Homer. Besides the enrichment score, each motif is characterized by a number of additional performance measures. First, the precision-recall curve that would be obtained if peak sequences were classified based on the number of sites for the motif is calculated and the area under the curve is reported (Fig. 4E). We see that motif denovo_WM_9 can identify about 40% of the peaks with high precision. Second, Crunch investigates to what extent the number of predicted binding sites in a peak correlates with the overall significance (*z*-value) of the peak (Fig. 4F). We see that as soon as the *z*-value is larger than about 8, peaks are virtually guaranteed to contain at least 1 site for denovo_WM_9. In addition, peaks with very high *z*-scores tend to have more than one binding site. Third, if the sites of the motif correspond to the locations where the immuno-precipitated protein associates to the DNA, we would expect that the sites would tend to occur near the center of the binding peak, i.e. where the ChIP coverage is highest (see also Suppl. Fig. S3). For each motif, Crunch also reports the distribution of coverages at binding sites and the distribution of coverages at all positions in the peak (Fig. 4G). We see that the ChIP coverage at binding sites for denovo_WM_9 is more than 9 times higher than at random positions in peaks. Finally, for each motif Crunch also reports the number of peaks for which binding sites are predicted to occur (Fig. 4A). Note that, while denovo_WM_9 occurs in about 50% of all peaks, its occurrence is highly specific, whereas the GFY-Staf motif occurs in almost all peaks but is much less specific. All other motifs in the set occur in only a small subset of the peaks.

All these results of the motif analysis are available through an interactive graphical interface that is automatically generated for each dataset that is submitted to Crunch, allowing the user to explore the results in detail. For example, a user can decide to select from all binding peaks only those peaks in which sites for a given motif are predicted to occur. In addition, to allow further downstream analysis of the binding site constellations in peaks, flat files are provided that annotate each peak with the precise occurrences of all the motifs from the complementary set {*w*}. PWM files for all the reported motifs are also provided.

### Biological interpretation of the BRCA1 results

Here we briefly discuss to what extent Crunch’s results on BRCA1 are consistent with what’s known in the literature. BRCA1 is a tumor suppressor that, since it was first uncovered in 1990 (Hall et al. 1990), has been subject to extensive studies. Mutated BRCA1, together with BRCA2, is reported to be responsible for approximately two thirds of all familial breast cancer cases, whereas about 10% of all breast cancer cases are familial (Deng and Brodie 2000). In addition, mutations within these genes increase risk for ovarian, pancreas, uterus, cervix and prostate cancers (Rosen et al. 2006). BRCA1 is involved in several different cellular pathways including DNA damage repair, cell cycle check points, centrosome duplication, transcriptional regulation as well as the immune response (Deng and Brodie 2000; Rosen et al. 2006). Although BRCA1 is a DNA-binding factor with a preference for binding at sites of damaged DNA, it does not bind DNA specifically in that it has a clear binding sequence motif (Rosen et al. 2006; Starita 2003). However, it interacts with a number of different proteins including sequence-specific DNA binding factors and RNA polymerase II. For example, in the context of DNA damage repair, BRCA1 builds a large complex called BASC (Wang et al. 2000).

Although the statistics in Fig. 4 suggest that the denovo_WM_9 motif best represents the binding preferences of BRCA1, it is known that BRCA1 does not bind DNA sequence-specifically and, therefore, denovo_WM_9 thus most likely represents the binding specificity of another TF that associates with BRCA1. Early studies found that a very similar motif occurs primarily in TATA-less promoters and is involved in the expression regulation of about 5% of all human genes, in particular genes involved in protein synthesis and cell cycle regulation, especially in the G1-to S-phase transition (Wyrwicz et al. 2007). Interestingly, BRCA1’s action in cell cycle regulation has also been reported to be involved in the same transition (Fabbro et al. 2004). Crunch finds that the closest known motif is a motif from Homer’s library that describes the binding specificity of the TF ZBTB33, also known as KAISO (Fig. 4D). KAISO has been associated with breast cancer and especially with BRCA1-related breast cancer (Vermeulen et al. 2012). We thus hypothesize that KAISO is a key interaction partner of BRCA1 and that the denovo_WM_9 motif describes the binding specificity of the KAISO TF. A similar hypothesis was put forward in (Wang et al. 2012).

Besides denovo_WM_9, Crunch finds five additional motifs that substantially increase the enrichment score and have binding sites within the binding peaks (Fig. 4A). The most significant of these motifs is a motif associated with the TF CREB3 as inferred from HT-SELEX experiments (Jolma et al. 2013) and Crunch reports that the CREB3 motif is highly similar to the motifs of the ATF and JUN family proteins (see Suppl. Figure S4). BRCA1 is well known to directly bind to CREB-binding protein (CBP) (Pao and Janknecht 2000) as well as to directly bind to ATF1 (Rosen et al. 2006) and JUN proteins (Hu and Li 2002). We thus hypothesize that a complex of BRCA1-CBP-CREB, a complex of BRCA1-ATF1 or a complex of BRCA1-JUN is binding to a subset of our BRCA1 binding peaks.

For the next two motifs, SPI1/PU1 and RFX3, we could not find any support in the literature. Although both together bind only 18 of our peaks, it might still be interesting to further investigate these two motifs in relation to BRCA1. The GFY-staf occurs in almost all peaks and has previously been associated with chromatin regions bound by H3K4me3 which are specifically open in breast cancer cells (Hong et al. 2012). Finally, we find the STAT1 motif, a TF that has also been reported to directly bind BRCA1 (Rosen et al. 2006). In summary, Crunch correctly identifies the key interaction partners of BRCA1 that have previously been described in the literature, and identifies two new motifs that appear to account for BRCA1 binding in a small subset of the peaks.

### Crunch’s noise model accurately reflects fluctuations in ChIP read densities

Assessing the accuracy of the predicted binding peaks of any ChIP-seq analysis tool is complicated because of the absence of an independent ground truth. However, since peak callers use a statistical model to distinguish true binding peaks from ‘random’ fluctuations in read density, the accuracy of peak finding relies on the accuracy of the statistical model. In the BRCA1 example above, we saw that the distribution of *z*-values inferred by our noise model accurately tracked the expected standard normal distribution, supporting that Crunch’s noise model correctly captures the statistics of ChIP-seq coverage fluctuations across the genome. We find that this is observed for most of the ENCODE ChIP-seq datasets.

For each of the 128 datasets, we quantified the difference between the expected distribution of *z*-values and the observed one by the root of the average squared-devation of the observed log-density of *z*-values and the standard-normal log-density, i.e. 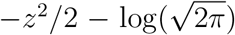. Because the observed distribution is expected to deviate from the expected standard normal at high *z*-values, this comparison was restricted to the range *z* ∈ [–5, 3], which covers almost 99.9% of all windows. As shown in Fig. 5 we observe a very good match between the expected and observed distribution of z-value for about 90% of all data-sets. Note that, even for for the worst cases, i.e. Fig. 5D, the observed distribution follows the standard normal over the range *z* ∈ [–2.5, 2.5] which includes almost 99% of all windows.

**Figure 5:**
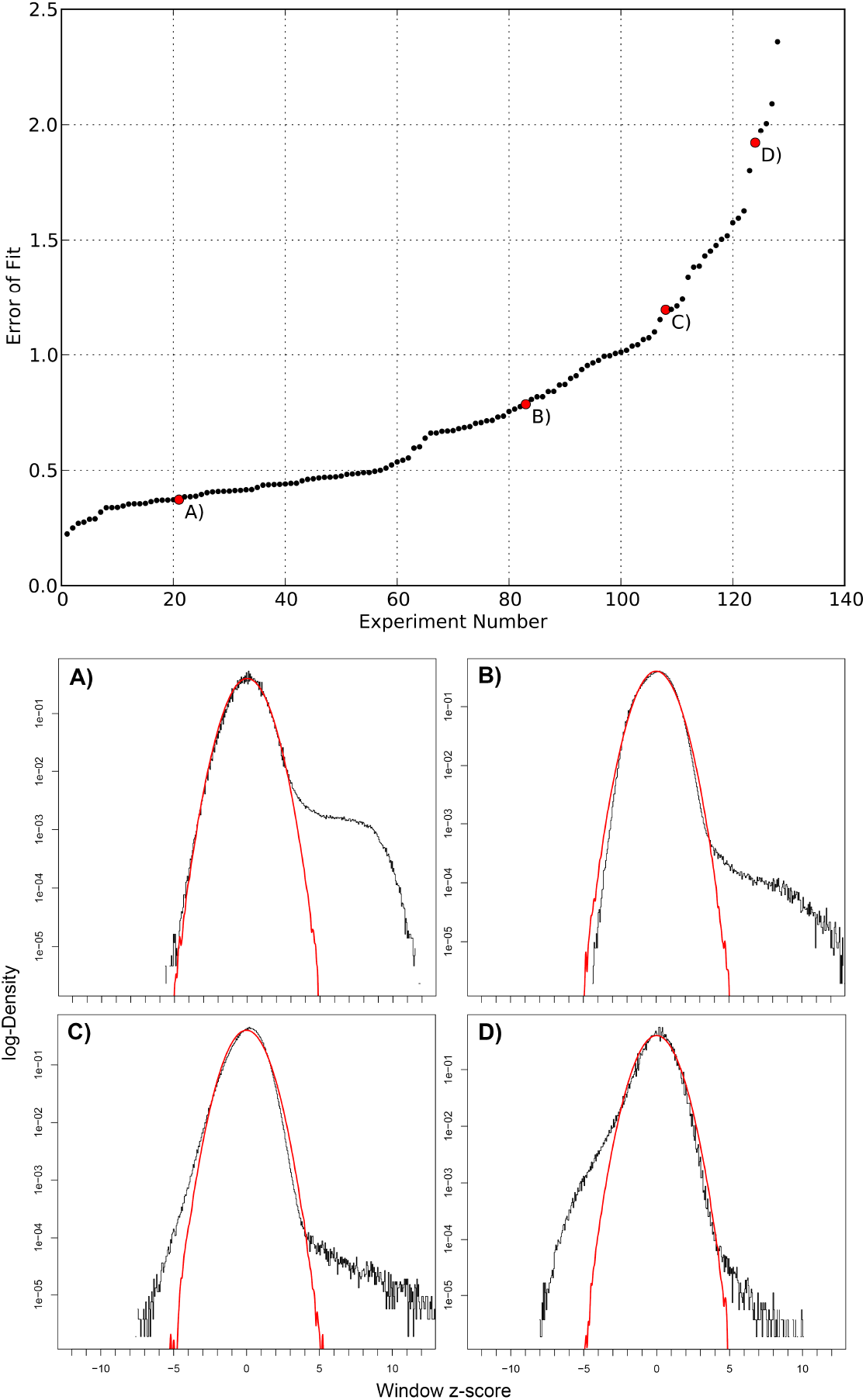
Genome-wide fluctuations in ChIP signal fit Crunch’s noise model. The top panel shows a cumulative distribution of the mean squared deviation of the assumed and observed *z*-value distribution across the 128 ENCODE ChIP-seq experiments. The bottom panels A-D show the observed (black) and expected (red) *z*-value distributions for four example datasets. The mean squared deviation for these 4 datasets is indicated in the top panel.

As far as we are aware, Crunch is the only peak finding tool for which the noise model is explicitly supported by the data. Furthermore, the good match between Crunch’s noise model and observed fluctuations in ChIP read densities provides confidence in the statistical significance that Crunch assigns to the enriched regions. Vice versa, because Crunch provides a figure with the comparison of the observed and expected *z*-value distribution in its report, users can immediately check whether their data shows an aberrant distribution of ChIP signal fluctuations.

### Crunch’s *de novo* motifs often outperform literature motifs

A second deliverable of ChIP-seq analysis is the analysis of sequence motifs occurring within the peaks. As discussed above, Crunch uses both a large library of known motifs, as well as motifs found *de novo* on the dataset in question, to determine a complementary set of motifs that jointly explain the observed binding peak sequences. To assess the importance and the quality of the motifs found de novo, we compared the enrichment scores of the top *de novo* and library motifs for each of the ENCODE ChIP-seq datasets (excluding 7 datasets for which less than 200 peaks were found). Note that, for most of the TFs that were immuno-precipitated in these datasets, the library of known motifs already includes several motifs representing these TFs, and it also includes all the ENCODE motifs that were previously inferred from these same datasets (Wang et al. 2012). We thus expected that, for most datasets, one of the many known motifs would outcompete the motifs that Crunch found *de novo.* In contrast, we find that for the majority of datasets, the best *de novo* motif outcompeted all known motifs (Fig. 6).

**Figure 6:**
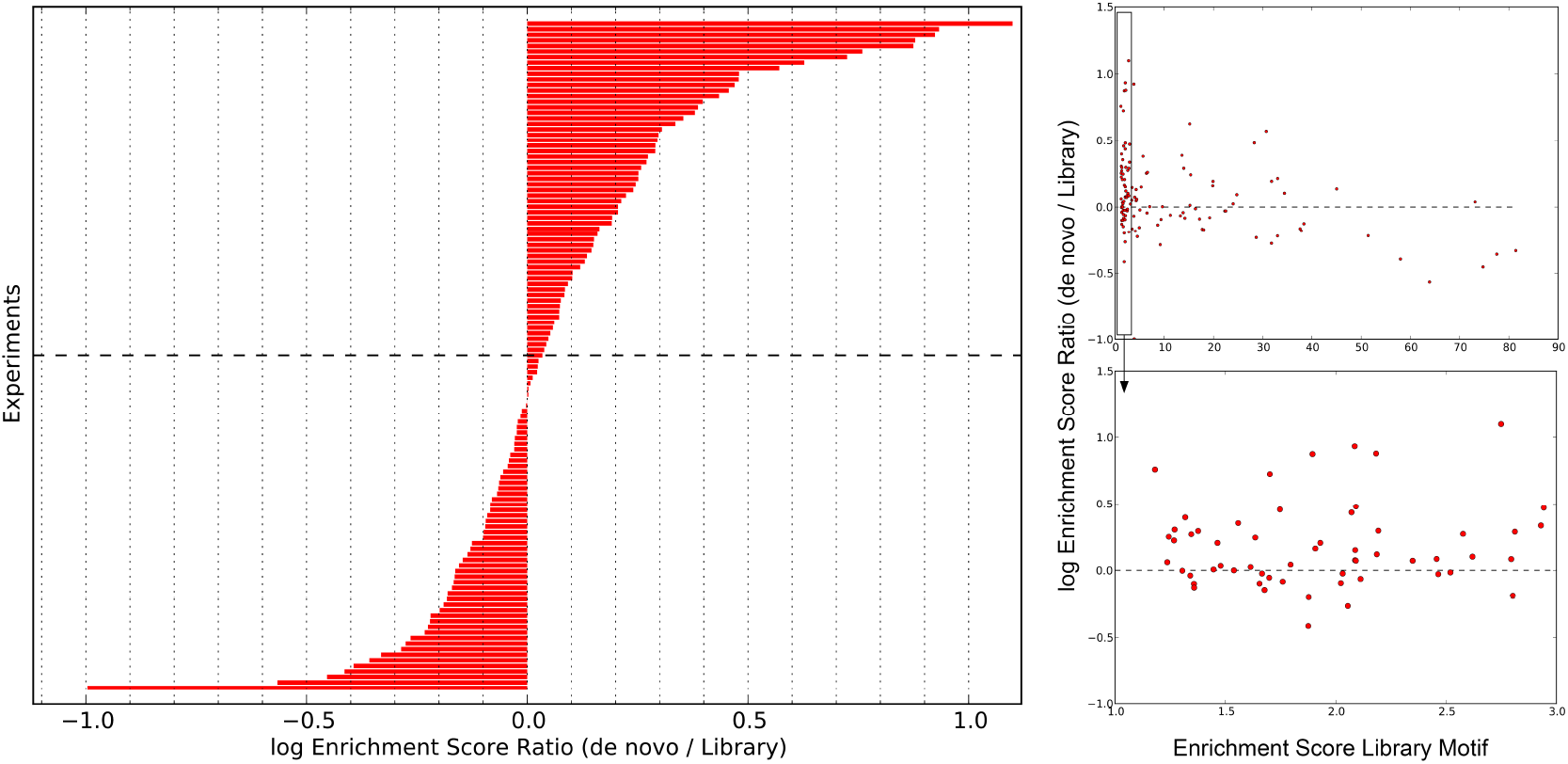
Comparison of the performance of known and *de novo* motifs. The left panel shows a bar plot of the differences in log-enrichment scores of the top motif found *de novo* by Crunch and the top motif from the library of known motifs across the 121 ENCODE datasets for which at least 200 binding peaks were identified. A positive difference means that Crunch’s *de novo* motif outperformed all library motifs. The panels on the right show the difference in log-enrichment between the best *de novo* and library motifs (vertical axis) as a function of the enrichment score of the best library motif. The bottom panel shows a zoom in of the boxed region on the left in the top panel.

Crunch not only provides a superior novel motif for the majority of the experiments, in many cases the novel motif performs substantially better than any existing motif. Note that, since there are typically 500 sequences in our test set, a difference in log-enrichment of 0.2 or more corresponds to likelihood ratio of e^0.2×500^ ≈ 2.7 × 10^43^ or larger, which is observed for more than a quarter of the datasets. Finally, we note that Crunch identifies superior *de novo* motifs especially when there was no high scoring motif in the library of known motifs, i.e. for more than two-thirds of the cases where the best known motif had an enrichment of less than 3, Crunch found a better motif *de novo* (Fig. 6, right panels).

### We observe two types of DNA-binding factors: solitary binders and co-binders

For TFs that specifically bind the DNA, one might expect that the binding peaks should in principle be explainable by just a single motif, i.e. the motif representing the sequence specificity of that TF. To quantify to what extent additional motifs beyond the top motif are needed to account for the observed peaks we defined, for each dataset, the ‘additional information’ as the difference in log enrichment of the entire motif set and the log enrichment of the top motif. As shown in Fig. 7 (left panel), the histogram of additional information across datasets appears multi-modal. First, there is a large peak of TFs that appear to be explained by a single motif, i.e. there is virtually no additional information. However, for the majority of TFs there is significant additional information in the other motifs, with the additional information exceeding 1.5 for some, corresponding to a 4.5 times higher enrichment. For TFs that were immuno-precipitated in two different cell lines, the additional information is generally very similar in both cell lines (Fig. 7, right panel), supporting that the additional information is an intrinsic property of the TF. In summary, while there is a minority of ‘solitary binding’ TFs for which the binding peaks are explained by a single motif, the binding of most TFs depends substantially on ‘co-binding’ of other factors, and the extent to which a TF acts in cooperation with other factors appears conserved across experiments and cell lines.

**Figure 7:**
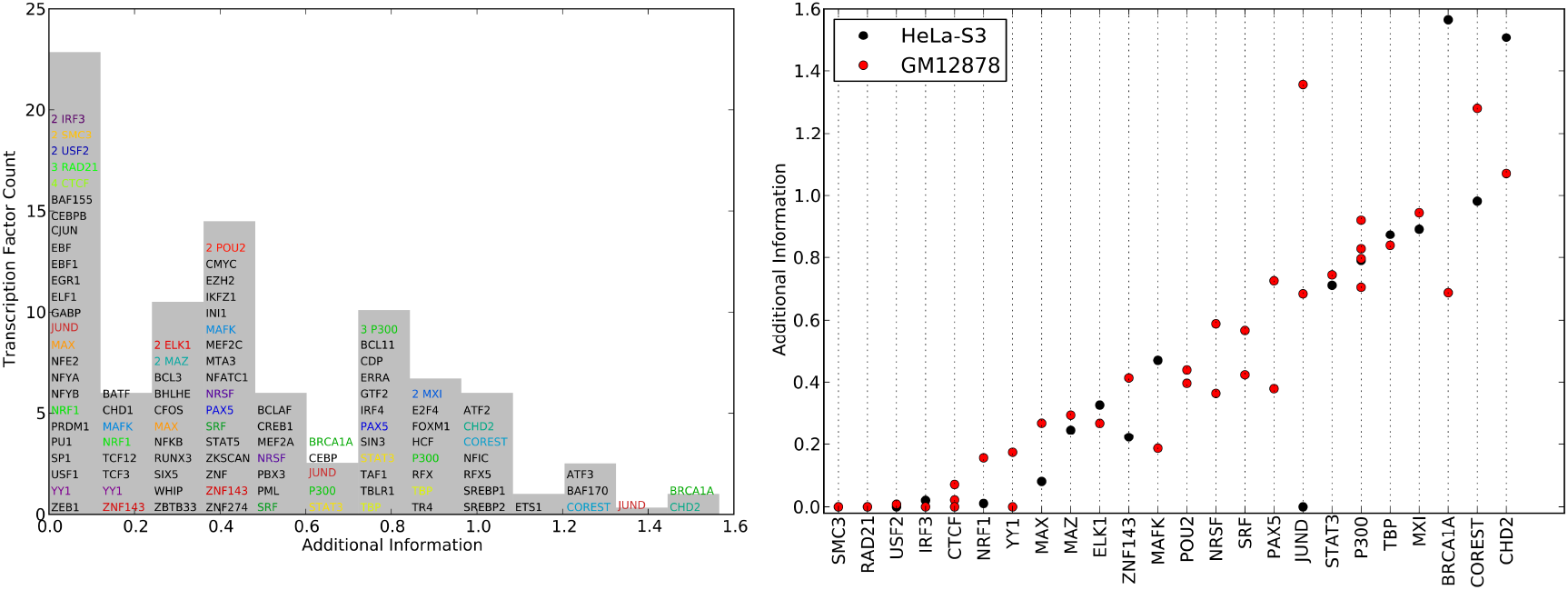
Contribution of motifs in the complementary motif set beyond the top motif. Left panel: Histogram of the additional information for all complementary motif sets from the ENCODE data. The height of the bin corresponds to the number of unique TFs in the bin. TFs with multiple experiments are shown in colored font, and whenever multiple experiments of the same TF occur in the same bin, the multiplicity is indicated by a number. Right panel: Additional information across multiple experiments and cell lines for the 24 TFs for which multiple experiments were performed. Each column corresponds to a TF and each dot to an experiment, with the cell line indicated by the color of the dot. The TFs are sorted by their mean additional information.

Inferred motifs are consistent across replicate experiments We next investigated to what extent the complementary motif set is consistent across different experiments with the same TF. We divided the set of TFs into ‘solitary binders’ and ‘co-binders’ depending on whether the average additional information in the complementary motif set was smaller or larger than 0.2. There were 16 co-binding TFs for which multiple experiments were performed. For each pair of experiments performed with the same co-binding TF we compared the motif sets using a consistency score which runs from 0, when there is no overlap between the motif sets, to 1, when the exact same motifs occur in the exact same order (see Methods). With the exception of a few motifs such as Stat3, P300, and CoREST, the large majority of co-binding TFs shows highly consistent motif sets across the experiments, including experiments on different cell lines (Fig. 8). This suggests that, for most co-binding TFs, the complementary motif set is an inherent characteristic of the TF. Moreover, it suggests that Crunch can successfully identify such co-binding motif sets.

**Figure 8:**
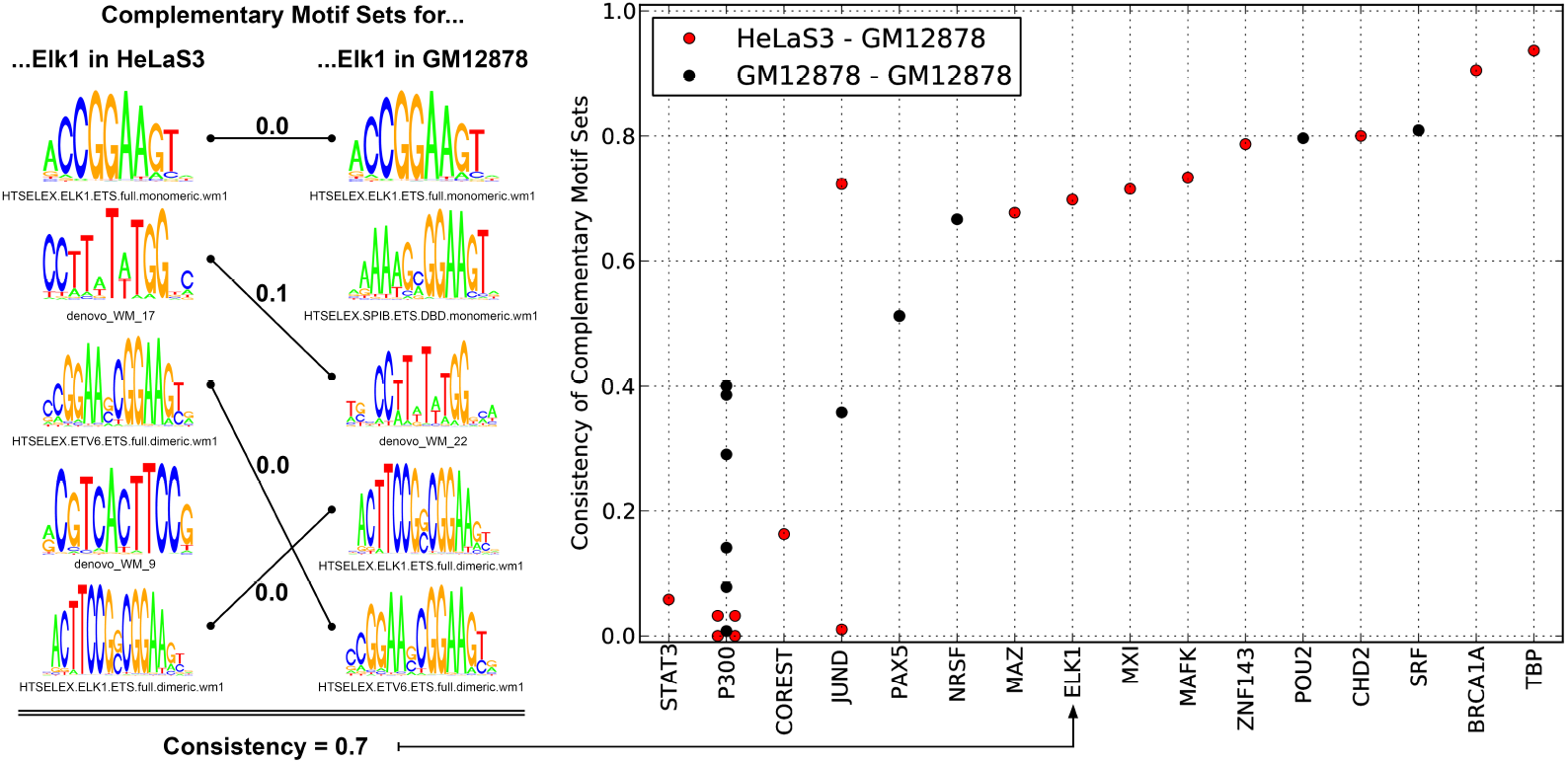
Consistency of the complementary motif sets of co-binding TFs across experiments. Left panel: Example of the complementary motif sets obtained from experiments with Elk1 in two different cell lines. Similar motifs are connected by lines with the number indicating the distance between the motifs (see Methods). Both sets contain another unpaired motif at position six which is not shown here. The consistency score for these two sets is 0.7 (see Methods). Right panel: The consistency scores for all pairs of experiments with co-binding TFs. Each column corresponds to a co-binding TF and each dot to a pair of experiments with that TF. The color indicates whether the experiments were performed in the same (black) or different (red) cell lines.

For solitary binding TFs the binding peaks are explained by a single motif and we next investigated to what extent these solitary motifs are consistent across experiments. There were 8 solitary binding TFs for which multiple experiments were performed, with 22 experinments performed in total with these factors. For each of these experiments, we extracted the top 5 known motifs with the highest enrichment scores. We then compared the sets of top 5 motifs across all pairs of experiments performed with the same TF using the same consistency score. Note that the consistency again runs from zero for disjoint sets to 1 for sets of identical motifs in the exact same order. We find that, of the 8 TFs tested, only the experiments on USF2 showed disagreement on the order of the top motifs. All others showed very high to perfect consistency, preserving the relative order even of highly similar motifs (Fig. 9). This suggests that binding specificity of the TF is independent of the details of the experiment and that, more importantly, Crunch’s results can be used to unambiguously select an optimal motif for solitary binding TFs.

**Figure 9:**
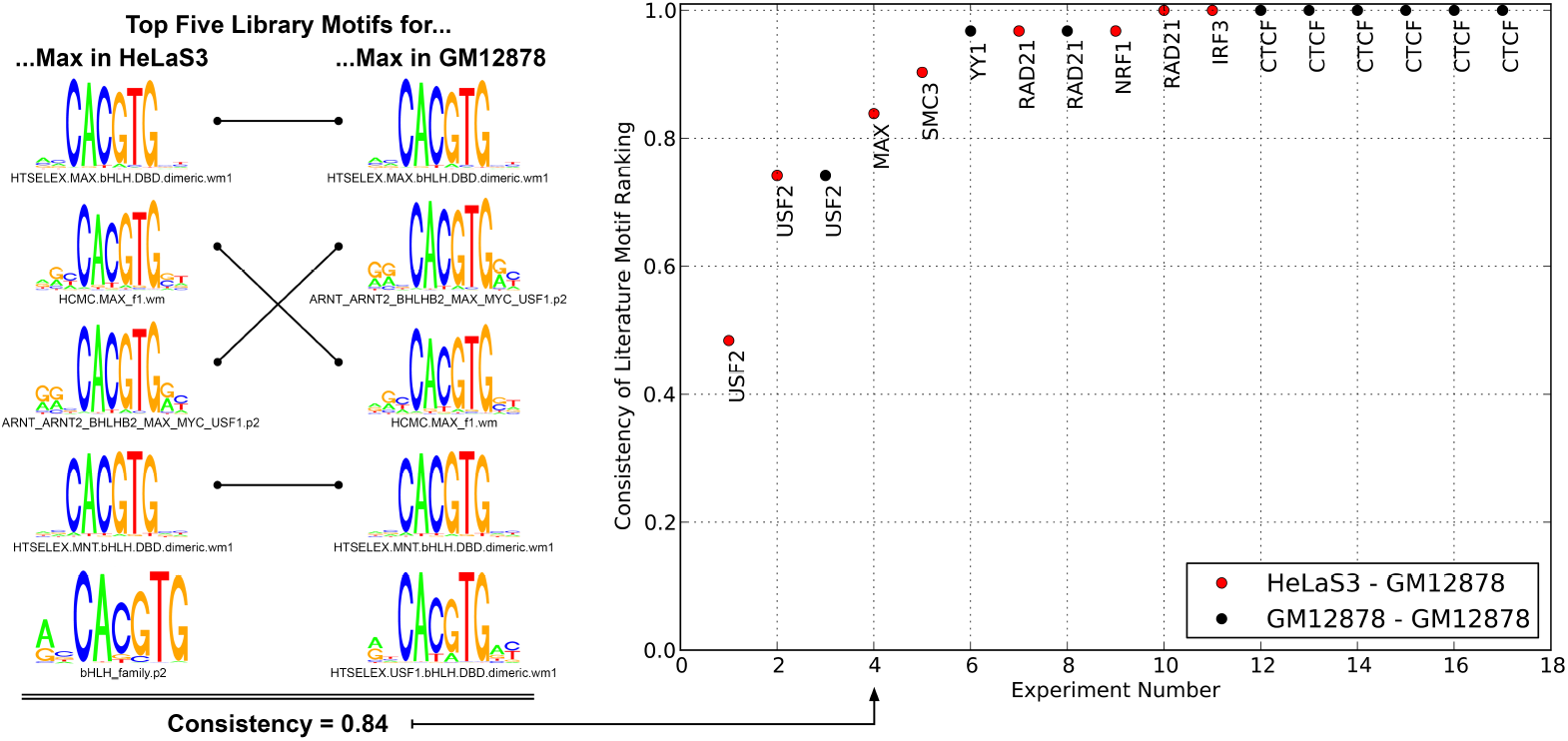
Consistency of the motifs of solitary binding TFs. Left panel: The top five library motifs for two experiments with the TF Max, in two different cell lines. Identical motifs are connected by lines. For this example, the consistency of the two sets of motifs is 0.84 (see Methods). Right panel: Consistencies of the top five library motifs for all 17 pairs of experiments performed with an identical solitary binding TFs. Each dot corresponds to an experiment pair, the name of the TF is indicated next to the dot, and the color indicates whether the pair of experiments was performed in the same (black) or different (red) cell lines.

## 5 Discussion

Although a large number of tools are available for performing the various steps of ChIP-seq analysis, in practice ChIP-seq analyses are very far from standardized, and the procedures used by different labs are so variable that it is virtually impossible to meaningfully compare ChIP-seq results from different studies. Moreover, implementing an accurate ChIP-seq anaysis pipeline requires in-depth expertise in selecting and combining an appropriate set of tools, tuning their parameters, and implementing necessary customizations. Consequently, experimental researchers wishing to perform ChIP-seq analysis are typically dependent on collaborations with computational experts.

To address these issues we developed Crunch, a completely automated pipeline that performs all ChIP-seq analysis steps in a standardized manner. Crunch does not require any computational expertise of the user. Raw data can simply be directly uploaded to the Crunch webserver, all analysis is performed automatically, and results are provided both through an interactive graphical web interface as well as through flat files for further downstream analysis. In this way, Crunch allows any user not only to perform ChIP-seq analysis on their own data, but also to systematically compare different ChIP-seq datasets. Moreover, Crunch provides extensive reports on all steps of the analysis that make it easy for users to diagnose potential problems.

Beyond providing a standardized pipeline, Crunch also introduces a number of novel features. In contrast to all peak finding tools that we are aware of, Crunch uses a noise model for fluctuations in ChIP read densities that is automatically fitted to the data, and that demonstrably matches the statistics of the fluctuations that are actually observed in the data. Probably the most important novel feature of Crunch is that a comprehensive regulatory motif analysis is integrated into the pipeline. Capatalizing on motif finding tools previously developed in our lab (Arnold et al. 2012; Siddharthan et al. 2005), Crunch uses a combination of *de novo* motif finding and prediction of binding sites for a large library of known motifs to explain the observed binding peak sequences in terms of the constellations of regulatory motifs that occur within them. In particular, Crunch automatically identifies a minimal set of complementary motifs that jointly best explain the observed peak sequences. Included among Crunch’s result files are files listing not only all binding peaks, their chromosomal locations, and ChIP signal strength, but also a comprehensive annotation of the regulatory motifs occurring within these sequences, as well as statistical analyses of the co-occurrence of all these motifs and their contribution to explaining the peak sequences. We believe that these results will be especially valuable for further downstream analysis of how sequence motif constellations determine the genome-wide binding patterns of TFs that are observed *in vivo.*

We demonstrated the power of the method by applying Crunch to a large collection of ChIP-seq datasets from the ENCODE project. First, we found that the motifs that Crunch identified *de novo* outcompeted all known motifs for the majority of the datasets, supporting the power of Crunch’s motif finding procedures. Second, Crunch’s analysis showed that immuno-precipitated factors can be naturally divided into ‘solitary binding TFs’ where the peak sequences are characterized by a single binding motifs, and ‘co-binders’ where the peak sequences contain sites for a combination of complementary motifs. We also found that, for most TFs, the set of co-binding motifs is consistent across different experiments and cell lines.

Although we have here focused on results of applying Crunch to data from ChIP-seq experiments with TFs, Crunch is not limited to such data. In particular, Crunch can also be applied to other types of data. For example, using data from DNase-seq experiments, Crunch can be used to identify regions of open chromatin. Similarly, using data from ChIP-seq experiments for the H3K4me1 or H3K27ac chromatin marks, Crunch can be used to identify the locations of cis-regulatory enhancers. In this context we believe that Crunch’s comprehensive motif analysis will be especially valuable for downstream analysis of how the genome-wide location of open chromatin and active enhancers is determined by local sequence motifs. In future studies we imagine that Crunch can be used to systematically investigate changes in genome-wide TF binding and open chromatin patterns can be understood in terms of local constellations of regulatory sites.

## 6 Methods

The Crunch analysis pipeline has been implemented using the Anduril workflow engine (Ovaska et al. 2010). Below we describe all analysis steps in the order in which they occur in the pipeline (see Fig. 1).

### Quality Control and Adapter Removal

Raw sequencing reads are typically provided in FASTQ format and Crunch directly takes such FASTQ files as input. These raw reads can have diverse quality due to uncertainty in base calling and errors in the sequencing itself, and they potentially contain artefacts such as sequenced parts of sequencing adapters. To avoid contaminating downstream analyses with low quality or erroneous sequence, we perform an initial quality filtering, followed by adapter removal, and a final round of quality filtering as follows.

In the first round of quality filtering we discard reads that are either shorter than 25 nucleotides (nt), contain more than 2 ambiguous nucleotides (N’s), or have an average Phred base calling quality score below 20 (corresponding to an error rate of 1%). As sequencing quality tends to decrease from the 5’ to the 3’ end of the read, we select the longest 5’ prefix of the read that has an average Phred score of at least 20. The chosen prefix is maintained if it has a minimum length of 25 nt.

In the adapter removal step, we focus exclusively on 3’ adapters, i.e. adapters that get (partially) sequenced if the sequence of interest (i.e. the fragment) is shorter than the length of the sequenced read. Crunch first aims to determine, for any given data set, which 3’ adapter was used. Crunch uses a list of known Illumina adapters (Illumina incorporated 2018), which can be extended with custom adapters by the user if desired. For each adapter in the list, prefixes of lengths 14, 16, 18 and 20 nt are mapped to 250’000 randomly chosen reads from the data set, allowing up to 2 mismatches. The adapter with the highest number of matches is chosen as the putative adapter sequence. This adapter sequence is then trimmed stringently from the reads according to the procedure previously described in (Stadler et al. 2011) and Suppl. text S2.

After adapter removal, we first remove all sequences whose remaining length is less than 14 and additionally remove low complexity reads. We define the complexity of a read as the entropy 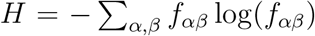, with *f_αβ_* the frequency of dinucleotide *αβ* in the read. All reads with *H* < 0.5 log(16) are removed (Stadler et al. 2011).

### Mapping

After filtering, Crunch maps the remaining reads to the reference genome using Bowtie version 1.1.1 (Langmead et al. 2009). Bowtie’s parameters are set such that for every read all mapping positions with the least number of mismatches get reported (-a – strata – best) allowing for at most three mismatches (-v 3) and skipping reads when the number of mapping positions exceeds 100 (-m 100). Multi-mapping reads are uniformly distributed over all mapping positions, i. e. each mapping of a read is assigned a weight equal to one over the number of mapping positions. We store these aligned reads in a BED-like format called BEDWEIGHT, which besides specifying the location of the mapping, also specifies the mapping’s weight. To allow visualization of the ChIP profiles in a genome browser, Crunch produces downloadable WIG files of the aligned reads at a 100 nt resolution.

### Fragment Size Estimation

After shearing and pulling down the DNA, ChIP-seq protocols for sequencing library preparation generally include a step that selects fragments in a certain size range. Since the lengths of the selected fragments are typically significantly longer than the length of the sequencing reads, reads are produced from both ends of each double-stranded fragment. Consequently, the read distribution in the neighbourhood of a protein binding site typically shows two peaks on opposite strands of the DNA, at a distance approximately equal to the typical fragment length (Schmid and Bucher 2007) so that the typical fragment length can be inferred from the correlated occurrence of reads on opposite strands at a particular distance *d*. Crunch estimates the fragment size by finding the distance *d* that maximizes the correlation function

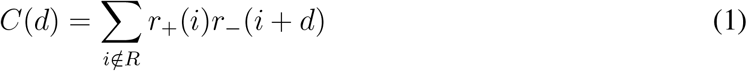

where *r*_+_(*n*) and *r*_(*n*) are indicator functions that equal one if a read occurs at position *n* of the, respectively, plus or minus strand, and zero if no read occurs. The sum in equation (1) is over all genomic positions excluding regions annotated as repeats (which we denoted by *R*). Note that we use indicator functions rather than raw read counts to avoid the correlation function *C*(*d*) to be dominated by a few positions positions with large read counts. *C*(*d*) is computed for *d* ranging from 0 to 600, and a typical resulting cross-correlation function is depicted in Fig. 2.

As has been observed previously, e.g. Landt et al. (2012), for some datasets a local maximum in *C*(*d*) occurs at *d* equal to the read length (see Suppl. Text S3). This is an artefact that we believe results from the ambiguous mapping of palindromic repeats. To avoid such artefactual fragment length estimates, we only consider local optima in *C*(*d*) for *d* larger than the read length.

### Peak calling: Identifying enriched regions

The first step in our peak calling procedure is to identify genomic locations that are enriched for fragments from the chromatin immuno-precipitation. Conceptually, if there is no binding of the immuno-precipitated protein at the locus, then the difference in ChIP and background fragment densities should result from random fluctuations, while when there is binding at the locus, then the ChIP fragment density will be higher than in the background sample. To this end we will have to compare, genome-wide, the fragment densities in the ChIP and background samples.

First, for each mapped read in both ChIP and background sample, we estimate the central position of the corresponding fragment to be half a fragment size toward the 3’ direction, i.e. forwards for reads on the plus strand and backwards for reads on the minus strand. Next, we count the number of ChIP fragments in sliding windows along the genome. By default Crunch uses sliding windows of length 500, shifting windows by 250 bps at a time. The choice of window length 500 is a tradeoff between obtaining sufficient mapped reads to measure local fragment density with reasonable accuracy, and obtaining sufficient spatial resolution to ensure that windows cover only one or a few binding peaks (a single binding peak’s width is roughly twice the fragment length, see below). If desired, the user can change the window length and the size of the shift between neighboring windows.

For each ChIP window, we count the number of reads in the background sample in a window of length 2000 (by default) centered on the same position as the ChIP window. Since the background fragments are approximately uniformly distributed in the background sample, the background density fluctuates more slowly than for the ChIP sample (which has sudden peaks) and the average fragment density is also much lower than in ChIP binding peaks, so that a larger window is required to accurately estimate the background density. The user can again change this default if desired. The result is two vectors of fragment counts *n* and *m* for the ChIP and corresponding background windows, where fragment counts of possible ChIP or background replicates are summed.

Although the background fragment counts should ideally be approximately constant, we have observed that there is a small fraction of windows (typically around 0.1%) that show much higher counts than all other windows (see Fig. 3A). Inspection of these regions show that they typically correspond to highly repetitive regions that align poorly with genomes of closely-related species. We suspect that the fragment counts are aberrantly high in these regions because these repeats are much more abundant in the genome used in the experiment than in the reference assembly, causing reads to pile up in these regions. These regions do not obey the statistics that are observed for the vast majority of the genome, and this leads to a high rate of false prediction of binding peaks in these regions. We thus developed a procedure to filter out these regions with unusually high background signal. Reasoning that the distribution of fragment counts for normal background signals should have a roughly exponential tail, we fit the tail of the reverse cumulative distribution of the background fragment counts to an exponential distribution and determine the point at which the observed distribution starts deviating more than exp(0.5) vertically from the fitted exponential tail (this point corresponds to the red line in Fig. 3A). All windows with counts above this cut-off are excluded from further analysis.

Next, in order to compare the fragment densities in ChIP and background samples, we need to normalize by the total fragment counts. We first add a pseudo-count of 0.5 to the fragment counts for each window and then determine the sums *N*, and *M*, of the fragment counts across all windows in the ChIP and background sample. In previous work (Balwierz et al. 2009) we have shown that the fluctuations in the number of next-generation sequencing reads across experiments can be well approximated by a combination of multiplicative (which may results from uncontrolled variations both in the biological state of the cells and variations in the process of preparing a sequence library from the sample) and Poisson sampling noise (from the sequencing itself), which leads to an approximately log-normal distribution of read-counts. In particular, if *f* is the true fraction of fragments deriving from a certain locus, the probability to obtain *n* reads at the locus when sequencing *N* reads in total is approximately given by

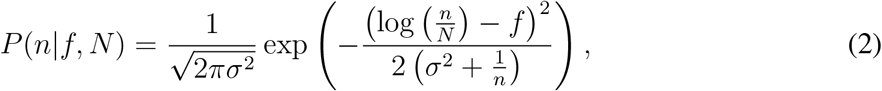

where *σ*^2^ is the variance of the multiplicative noise component, and the term 1/*n* corresponds to the variance due to the Poisson sampling noise (see (Balwierz et al. 2009) for details). If a given window corresponds to a region of the genome that was unbound, then the true fraction *f* should be the same in the ChIP and background sample. Consequently, the probability to obtain *n* out of *N* fragments in the ChIP sample when *m* out of *M* fragments were observed in the background sample, is given by:

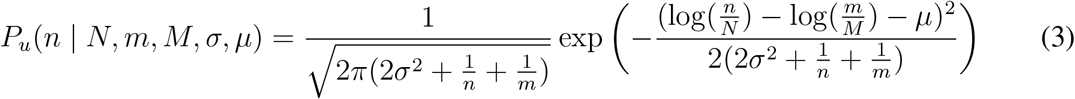

The term 2*σ*^2^ is the variance of the multiplicative noise component and the term 1/*n* + 1/*m* constitutes the contribution to the variance from the Poisson noise components of both the foreground and background samples. As a significant fraction of the reads in the foreground sample derives from bound regions, the fragment density in regions without binding is systematically lower in the ChIP sample than in the background sample. The parameter *μ* corresponds to the resulting overall shift in log-density in the unbound regions. Note that the distribution of equation (3) differs from the negative binomial distribution that is often used to model the statistics of next-generation sequencing read counts (Anders and Huber 2010). A negative binomial assumes that, instead of log-normal, the biological replicate noise is Gamma distributed.

If a window corresponds to a region that had binding of the immuno-precipitated factor, then the fragment density can be arbitrarily higher in the ChIP sample than in the background sample. Instead of making specific assumptions about this distribution, we will describe the probability *P_b_*(*n|N, m, M*) to obtain *n* out of *N* reads in the ChIP sample given *m* out of *M* by a uniform distribution in the difference *δ* of log-densities between ChIP and background. That is, let *δ* = log(*n/N*) – log(*m/M*), and let *W* = *δ*_max_ – *δ*_min_ correspond to the observed range in *δ* values across all windows. We then assume that, for a bound region, the *δ* value can take on any value in this range

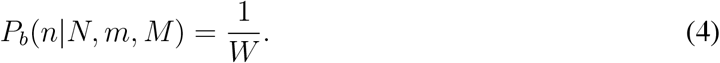

To model the joint distribution of both bound and unbound windows, we now assume a mixture model. That is, the overall probability for a window to obtain *n* out of *N* fragments in the ChIP sample given that it had *m* out of *M* fragments in the background sample is given by

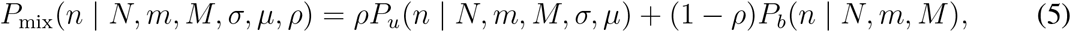

where *ρ* is the fraction of windows that are unbound, and *μ* and *σ* are the parameters of the noise model for unbound regions as described above. We then fit the parameters *μ*, *σ* and *ρ* by maximising the log-likelihood across all windows using expectation maximisation (see Suppl. Text S4 for details). Finally, since the Gaussian approximation in equation (3) becomes inaccurate when the raw fragment counts are only zero or one, we exclude windows where the ChIP fragment count is below 2.

After the parameters *μ*, *σ* and *ρ* have been fit, we compute a *z*-value for every window:

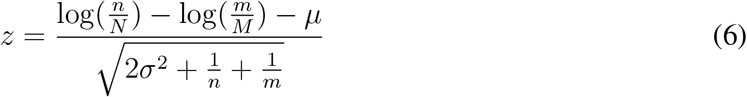

Note that, if there was no binding at any of the windows in the genome, the *z*-values should follow a standard normal distribution. As illustrated in Figs. 3B and 5, for most data-sets we find that the *z*-values of almost all windows indeed accurately follow a standard normal distribution, indicating that our statistical model successfully captures the noise distribution in the unbound regions, which constitute the vast majority of the genome. We also typically find that, as expected, a small fraction of windows shows substantially higher *z*-values.

As a final list of bound windows we select all windows with *z*-value over a threshold *z*_*_. Since the mixture model assigns a posterior probability to each window to be bound, we can easily estimate, for a given value *z*_*_, what the expected false discovery rate (FDR) is, i.e. the fraction of all windows with *z*-values larger than *z*_*_ which do not derive from the bound distribution *P_b_* (see Suppl. Text S5 for details). We set *z*_*_ by default such that the FDR is 0.1 (Fig. 3C), but this value can also be altered by the user if desired. Finally, since we chose the sliding windows to overlap, we merge overlapping windows that passed the threshold into larger bound regions.

### Peak calling: Identifying individual binding peaks within enriched regions

The enriched genomic regions that result from the analysis in the previous section are typically 500-1000 base pairs in length, which is significantly longer than the length of individual protein binding sites on the DNA. In the second step of peak calling we search for individual binding events by inspecting the ChIP signal within the regions at single basepair resolution. For each position in each significantly enriched region, we compute the number of foreground fragments that overlap it. Here fragments are reads that were extended from their 5’ end to fragment size in 3’ direction. The result is a ChIP coverage profile for each significantly enriched region (Fig. 3D).

To detect individual binding events we now fit the coverage profile of each enriched region to a mixture of Gaussian peaks plus a uniform background distribution. Approximating the data as if the coverage *C(i)* at each position *i* in the region were an independent observation, the likelihood of the mixture model takes on the following form:

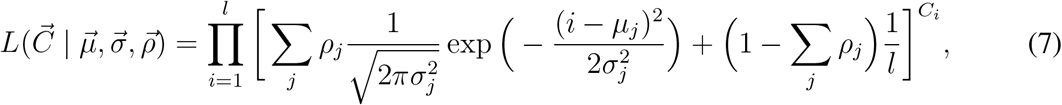

where *i* runs over all positions from 1 to *l* in the region, *C(i)* is the coverage at position *i*, i.e. the number of fragments that are overlapping position *i, j* runs over all Gaussian peaks in the model, *μj* and *σ_j_* are the central position and width of the Gaussian *j, ρj* is the fraction of all observations that belong to Gaussian *j*, and the last term corresponds to the uniform background distribution that accounts for the coverage not associated with the Gaussian peaks. Importantly, for individual binding sites on the genome, the width of the resulting coverage peak is a relatively well-defined function of the fragment length and we use this to constrain the widths *σ_j_* of the Gaussian peaks to fall within a range that is consistent with these peaks corresponding to single binding sites on the genome (Suppl. text S1 and Suppl. Fig. S5). We also use our knowledge of the typical width of individual binding peaks to constrain the numbers of Gaussians in the mixture model, i.e. we use as many Gaussians as can be fitted within the region given the expected widths of the Gaussians, with a minimum of 2 Gaussians. We again fit the parameters of the model by maximizing the likelihood of the coverage profile using expectation maximisation (Suppl. Text S6).

The optimization of the Gaussian mixture may result in some Gaussian components that highly overlap. We merge overlapping Gaussians if the difference between their mean positions is less than the sum of their standard-deviations, which roughly corresponds to the condition that there is no local minimum in the coverage profile between the two peaks. The *ρ*-weighted averages of the means and standard-deviations of the overlapping Gaussians are used to define the *ρ*, mean and standard-deviaton of the single merged Gaussian. Finally, for each Gaussian component, the binding peak is defined as the region [*μ* – *σ, μ* + *σ*] (see colored rectangular regions in Fig. 3D).

Note that, because the number of Gaussians used in the mixture is chosen to be an upper bound on the number of real binding peaks, some of the resulting Gaussian peaks may not exhibit a significant enrichment over background. We thus calculate a *z*-value for each individual peak and only retain those peaks with a *z*-value above the threshold *z*_*_ computed in the preceding section. The *z*-value is computed according to equation (6) using newly computed counts n which result from summing the contribution of the Gaussian peak in equation (7), together with the uniform background. The renormalized number of reads associated with peak *j*, in a region of length *L*, is given by

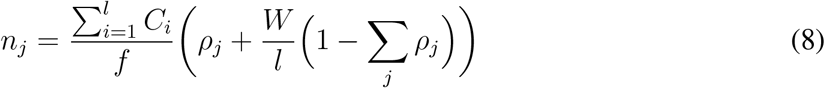

where *f* is the fragment size and *W* is the window length (500 by default). Note that the first factor corresponds to the total number of fragments assigned to the enriched region. The term proportional to *ρ_j_* corresponds to the fraction of fragments directly assigned to the peak by the mixture model, and the second term corresponds to the total number of background fragments in a region of length *W*.

### Association of binding peaks with genes and promoters

To annotate which genes may be regulated by the regulatory elements within the peak, we use our curated collection of promoters from SwissRegulon (Balwierz et al. 2009; Pachkov et al. 2013) and record the three closest promoters up- and downstream from the peak, as well as the genes associated with these promoters. In addition, for each peak we provide a link to the SwissRegulon genome browser, displaying the peak within its genomic context, including annotations of known transcripts, known promoters, and predicted transcription factor binding sites within these promoters.

### Regulatory Motif Analysis

Crunch performs an extensive regulatory motif analysis which aims to optimally explain the observed ChIP-seq signal in terms of the occurrences of regulatory sites for both novel and known regulatory motifs. To comprehensively characterize the regulatory sites occurring within each binding peak, Crunch uses a combination of *de novo* motif finding and binding site prediction for a large collection of known regulatory motifs. Using these predictions Crunch then finds a complementary set of regulatory motifs that optimally explains the observed ChIP-seq data using a novel approach that computes a likelihood of a given motif set under an idealized model of the chromatin immuno-precipitation process. A greedy optimization algorithm is used to find a set of complementary motifs that maximizes this likelihood. In addition, Crunch computes a number of different statistics to characterize the distribution of binding sites within peaks for all motifs within the complementary set.

To perform the motif analysis Crunch sorts the binding peaks by z-value and collects either all binding peaks, or the top 1000 peaks when there are more than 1000 significant peaks. The peaks are then randomly divided into two equally sized subsets: a training set {*P*_training_} that Crunch uses to find and optimize motifs, and a test set {*P*_test_} that Crunch uses to assess the performance of motif sets.

### *De Novo* Motif Finding

To identify novel motifs Crunch uses PhyloGibbs (Siddharthan et al. 2005), which implements a Bayesian model for assigning posterior probabilities to configurations of putative sites for a number of unknown regulatory motifs (with both the total number of sites and maximum number of different motifs defined by the user), and samples configurations in proportion to their likelihood using Markov Chain Monte Carlo. PhyloGibbs was specifically constructed to incorporate information from conservation patterns across orthologous genomic regions given their phylogenetic relationships. For each binding peak Crunch uses UCSCs pairwise genome alignments to extract orthologous sequences from related genomes, i.e. (hg19, mm9, rheMac2, canFam2, bosTau6, equCab2, monDom5) when using data from human or mouse, and (droSim1, droYak2, droEre2, droAna3, dp4, droWil1, droVir3, droMoj3, droGri2) when using data from Drosophila. These orthologous regions are then multiply aligned using T-Coffee (Notredame et al. 2000), as described in (Arnold et al. 2012). To enable the detection of several, potentially non-redundant, motifs we run PhyloGibbs six times with different settings: Either using phylogenetic information and multiple alignments or using only the sequence from the reference species, and searching for motifs of lengths of either 10, 15, or 20 nucleotides. In each case we are searching for two motifs simultaneously (-z 2) defining that both together are expected to have 350 binding sites within the 500 peaks of the training set. Further, we use a first order background model (-N 1). This procedure yields 12 predicted motifs, represented by position specific weight matrices (PWMs). Crunch then refines these PWMs using the MotEvo algorithm (Arnold et al. 2012). MotEvo uses an expectation maximization procedure to optimize a set of PWMs so as to maximize the likelihood of the input sequences as a mixture of PWM sites and background. Applying this procedure separately to each PWM using the sequences from the training set yields 12 refined motifs. Finally, since these procedures identify motifs of a predefined width, one often observes a core motif flanked by uninformative columns, i.e. columns with nucleotide frequencies matching the background frequencies. Crunch trims all 24 motifs from both ends until a column with information content of at least 0.25 bits appears. Thus, at the end of these procedures, we have at most 24 candidate *de novo* motifs that we will subject to further analysis.

### Library of known motifs

We have collected a large library of known mammalian regulatory motifs from the literature. This library consists of the motif libraries from JASPAR (Bryne et al. 2008), HOCOMOCO (Kulakovskiy et al. 2013), HOMER (Heinz et al. 2010), UNIPROBE (Newburger and Bulyk 2009), ENCODE (Wang et al. 2012), HT-SELEX (Jolma et al. 2013), and SwissRegulon (Pachkov et al. 2013), containing a total of 2325 PWM motifs. For each dataset, we fuse this library of known motifs with the motifs that were found *de novo* for that dataset to form a set of candidate motifs that we denote {*W*_lib_}.

### The enrichment performance score for a set of motifs

Crunch finds a set of complementary regulatory motifs {*w*} that can jointly ‘explain’ the observed peak sequences using an idealized representation of a ChIP-seq experiment. First, we approximate the genome by a pool of sequences *P* consisting of the observed binding peaks *P_o_* together with a very large set of ‘background’ sequences *P_b_* that have the same lengths and nucleotide composition, but are otherwise random. Second, we assume that, when immuno-precipitating with a protein *X*, the probability of detecting a particular peak sequence *p* as a peak is proportional to the average number *n_p_* of copies of *X* that are bound to the sequence (averaged over many cells). Our probabilistic model idealizes the ChIP-seq by assuming the probality of observing the set of peak sequence *P_o_* is given by the probability of sampling these sequences from the large pool *P*, and none of the other sequences, i.e.

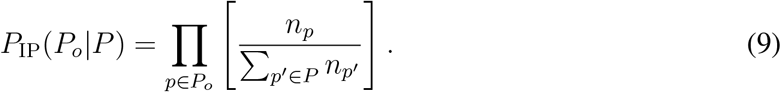

In order to relate the binding of *X* to the sequence of peak *p* we assume the following model. We assume that there is a set of motifs {*w*} representing the sequence-specificities of both *X* itself (if *X* is a sequence-dependent DNA-binding factor) as well as all other DNA-binding factors to which *X* binds either directly or indirectly, and that the total binding of *X* to sequence *ρ* is proportional to the total number of binding sites in *p* for the motifs in {*w*}. That is, we assume that the probability of observing sequence *p* as a peak sequence is given by

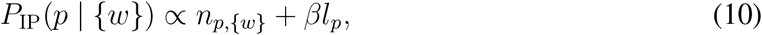

here *n*_*p*,{*w*}_ is the total number of binding sites for motifs of the set {*w*} within *p, l_p_* is the peak’s length and *β* corresponds to the amount of non-specific binding per nucleotide. We added a nonspecific binding term to the model because it strongly improves the performance of the model and it is well-known that TFs associate non-specifically with the DNA (mostly through electrostatic attraction). Using equation (10), we find that the probability of the set of observed peaks *P_o_* is given by

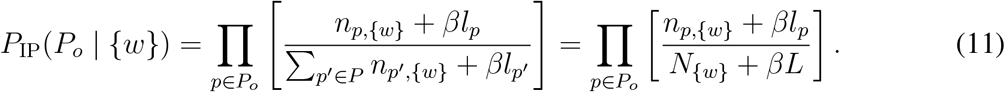

Where *N*{*w*} is the total number of binding sites for the motifs {*w*} within the large pool *P* and *L* is the total length of all sequences in the pool *P*. To assess the performance of the set of motifs {*w*} relative to random expectation, we use the difference between the log-likelihoods of observing the set *P_o_* given motifs {*w*} and when randomly sampling peak sequences. This log-likelihood difference is given by

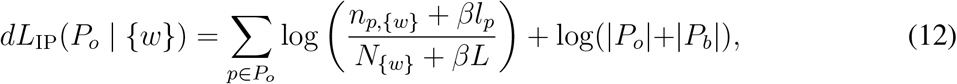

where |*P_o_*| and |*P_b_*| are the number of sequences in the sets of observed peaks *P_o_* and background sequences *P_b_*, respectively. Finally, taking the limit of assuming the background set *P_b_* much larger than the set of observed peaks *P_o_*, we find

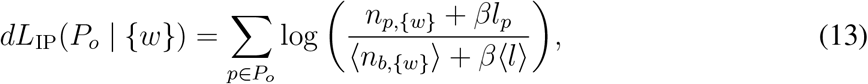

with 〈*n_b_*,{*w*}〉 denoting the average numbers of sites for motifs {*w*} per background sequence, and 〈*l*〉 the average length of the background sequences. The expression (13) gives the log-likelihood ratio of immuno-precipitating the true peak sequences *P*o** from a very large pool of sequences of equal composition and length, between a model in which sequences are sampled proportional to the number of sites they contain for motifs from {*w*}, and a model in which sequences are sampled randomly. To give a more intuitive measure, we transform this log-likelihood ratio into an ‘enrichment’ score as follows:

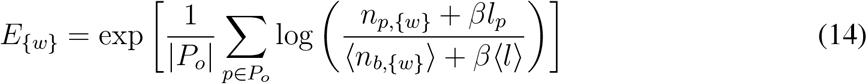

The enrichment *E*_{*w*}_ has a simple interpretation: it measures how much more likely it is (on average) to immuno-precipitate a true binding peak as opposed to a background sequence. Obviously, this quantity will depend on how we predict binding sites for {*w*} as well as on the choice of the parameter *β*. In the next section we discuss how Crunch determines these. Finally, we note that we denote the enrichment of a single motif *w* by *E_w_*.

### Binding site prediction and accounting for non-specific binding

In order to calculate the enrichment score for a set of motifs, we need to determine the number of binding sites *n_p,{w}_* for the peak sequences as well as for a large pool of background sequences. Although equation (14) effectively assumes an infinitely large pool of sequences, the score depends only on the average number of binding sites 〈*n*_*b*,{*w*}_〉 per background sequence, so that the enrichment score can be well estimated provided that the set of background sequences is large enough. We thus created a set of background sequences with 10 times as many sequences as the number of peak sequences. The background sequences are constructed to have the same distribution of lengths and nucleotide composition as the peak sequences. Since, as detailed below, the number of predicted binding sites *n_p,{w}_* depends on the parameter settings of the binding site prediction algorithm, and the enrichment score additionally depends on the non-specific binding parameter *β*, we divide the peak and background sequences into two equally sized subsets, *P*_1_ and *P*_2_, where the first ‘training’ set *P*_1_, containing {*P*_training_}, is used to optimize *β* and the parameters of the motif finding, and the second ‘test’ set *P*_2_, containing {*P*_test_}, is then used to calculate a final enrichment score *E*_{*w*}_.

We use the MotEvo algorithm (Arnold et al. 2012) to calculate the number of binding sites *n_p,{w}_* in each sequence *p* of *P*_1_ MotEvo is a Bayesian algorithm that models the input sequences as a mixture of non-overlapping sites for the motifs from {*w*} and nucleotides deriving from a background model (van Nimwegen 2007) and calculates posterior probabilities of binding sites to occur for each of the motifs in {*w*} at each of the positions in the sequences. MotEvo’s TFBS predictions depend on a set of prior probabilities {*π*}, with *π_w_* denoting the prior probability that a randomly chosen position in the input sequences corresponds to the start of a binding site for motif *w* ∈ {*w*}. Crunch runs MotEvo on the training pool *P*_1_ using a mode in which the parameters {*π*} are optimized so as to maximize the likelihood of the mixture model on the input data *P*_1_. Once the optimal parameters {*π*} are determined, Crunch additionally optimizes the non-specific binding parameter *β* so as to maximize the enrichment (14) for the training set *P*_1_ The optimal priors {*π*^*^} and optimal *β*_*_ are then fixed, MotEvo is run with these parameters on the test set *P*_2_, and a final enrichment *E*_{*w*}_ is calculated based on the binding site predictions on the test set.

It is worthwhile to note that the algorithm that MotEvo employs is equivalent to a thermodynamic biophysical model in which the priors {*π*} correspond to the concentrations of the TFs associated with the motifs in {*w*} and the posterior probabilities of the sites correspond to the fraction of time the sequence are bound by the respective TFs. In this interpretation the maximization of the priors {*π*} corresponds to maximizing the total binding free energy of the input sequences.

Another important point to note is that, as MotEvo only considers non-overlapping configurations of binding sites, redundant motifs compete for binding and consequently will not increase free energy of binding when added to {*w*}. More precisely, the sum of the optimized priors of two redundant motifs will be approximately equal to the optimized prior of one of the two redundant motifs by itself. In this way, addition of redundant motifs to the set {*w*} will generally leave the enrichment *E_{w}_* unchanged.

### Finding an Optimal Complementary Set of Motifs

Crunch searches for a minimal motif set {*w*} that maximizes the enrichment *E_{w}_*. As a complete search across all subsets of {*W*_lib_} is computationally infeasible, we use a greedy algorithm that maximizes *E_{w}_* by adding one motif at a time (Fig. 4A). Crunch starts by calculating the enrichment Ew for each motif in the library and sorts all motifs by this score. Since the motif library {*W*_lib_} is highly redundant, we typically find that any high scoring motif w on the list is accompanied by a number of highly similar but lower scoring motifs further down the list. These motifs are highly unlikely to end up in the final set {*w*} and for computational efficiency we remove these motifs from the list. That is, for any motif on the sorted list, all motifs that are highly similar but lower on the list are removed (see Suppl. Text S7 for details). We denote the sorted list of remaining motifs by {*W*_reduced_}. We initiate the motif set {*w*} with the top motif *w*_top_, i.e. the motif with maximal enrichment *E_w_*, and iterate:

1. For every motif *w* left in {*W*_reduced_} compute *E*_{*w*}∪*w*_.
2. Denote the motif *w* with maximal *E*_{*w*}∪*w*_ by *w*_*_.
3. If *E*_{*w*}∪w_*__ increases *E_{w}_* by more than 5%, add *w*_*_ to {*w*} and go to step 1. Otherwise, terminate the algorithm.

The cut-off of at least a 5 percent increase for each added motif was chosen so as to allow even motifs that add relatively little to be incorporated, while at the same time avoiding adding redundant motifs.

### Additional motif statistics

Besides the enrichment score *E_w_* we use a number of additional statistics to characterize the way in which each motif from the set {*w*} associates with the binding peaks (Fig. 4). First, for each motif Crunch reports what fraction of the binding peaks contains at least one site for the motif. Second, although Crunch uses the enrichment score *E_w_* to quantify the ability of the motif to explain the observed peaks, it also provides a standard precision-recall curve that shows how well binding peaks can be distinguished from background sequences based on the number of predicted sites (Fig. 4E). That is, by varying a cut-off on the total number of binding sites *T*, Crunch calculates what fraction of binding peaks have a number of sites larger than *T* (sensitivity) and what fraction of all sequences with more than *T* sites are true binding peaks (precision). The precision-recall curve shows the precision as a function of the sensitivity and the overall quality of the classification is quantified by the area under the curve, which is 1.0 for a perfect classifier and 0.09 for a random classification (because the true binding peaks are 10 times rarer than background sequences).

As a third measure, Crunch investigates to what extent the number of predicted binding sites in a peak correlates with the strength of the peak’s ChIP signal, i.e. whether sequences with more sites lead to more enriched peaks. Crunch both provides a graph showing a box plot of the distribution of the number of predicted sites as a function of ChIP signal strength (Fig. 4F), and calculates the overall Pearson correlation between the number of binding sites *n_p,w_* and the peak’s *z*-value. Finally, if binding sites for the motif were directly responsible for the immuno-precipitation of the fragments, then we would expect the positions of the binding sites within the peak region to co-localize with the peak of the ChIP signal. Crunch’s results also include a figure that shows the distribution of ChIP signal at predicted binding sites and, for reference, at all positions in the peaks (Fig. 4G). To quantify the enrichment of the ChIP signal at predicted binding sites we calculate a ChIP signal enrichment:

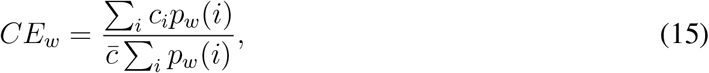

where *i* runs over all positions at which a binding site for motif *w* is predicted, *p_w_* (*i*) is the posterior probability (as assigned by MotEvo) for the site at *i, c_i_* is the ChIP signal (fragment coverage) at position *i*, and 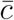 is the average ChIP signal in the binding peaks. Only binding sites with posterior *p_w_* (*i*) > 0.2 are included in this calculation.

Crunch also reports statistics on the co-occurrence of motifs from the set {*w*} within binding peaks. For each pair of motifs (*w,w*′) in {*w*}, Crunch calculates the Pearson correlation in the number of binding sites *n_p,w_* and *n_p,w_*′ across the binding peaks *p*. To visualize these correlations Crunch provides a heat map of between motif correlation over peaks (Fig. 4C).

### Consistency of Motif Sets

To compute the consistency *C* of two sets of motifs *S* and *S*′, we use the following measure, which is an extension of the Dice set similarity measure for ordered sets (Egghe and Michel 2003):

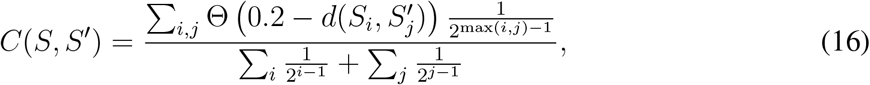

where *i* and *j* run from 1 to the number of motifs in *S* and *S*’, respectively, Θ(*x*) is the Heaviside step-function which is 0 if its argument is negative and 1 otherwise, and 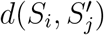 is the distance between motif *i* in *S* and motif *j* in *S*′ (Suppl. Text S7). That is, if the distance between two motifs is lower than 0.2, the motifs are considered to match (See Suppl. Fig. S6 for an example of motif distances among motifs). Note that the consistency *C*(*S, S*′) runs from 0 (no matching members) to 1 (two identical sets in the same order).

## Acknowledgments

We thank Bart Deplancke, Eileen Furlong, Felix Naef, Uwe Ohler, Jacques Rouge-mont, Michael Stadler, and Martin Vingron for valuable feedback on the ChIP-seq analysis procedures that are implemented in their labs.

## Correspondence

Correspondence and requests for materials should be addressed to Erik van Nimwegen.

